# Physiological and transcriptomic response of *Medicago truncatula* to colonization with high and low benefit arbuscular mycorrhizal fungi

**DOI:** 10.1101/2020.12.11.421693

**Authors:** Kevin R. Cope, Arjun Kafle, Jaya K. Yakha, Philip E. Pfeffer, Gary D. Strahan, Kevin Garcia, Senthil Subramanian, Heike Bücking

## Abstract

Arbuscular mycorrhizal (AM) fungi form a root endosymbiosis with many agronomically important crop species and both enhance the ability of their host to obtain nutrients from the soil and increase host tolerance to biotic and abiotic stressors. However, AM fungal species differ in the benefits they provide to their host plants. Here, we examined the putative molecular mechanisms involved in the regulation of the physiological response of *Medicago truncatula* to either *Rhizophagus irregularis or Glomus aggregatum,* a high or a low benefit AM fungus, respectively. Colonization with *R. irregularis* led to higher growth and nutrient uptake benefits than the colonization with *G. aggregatum*. These benefits were linked to an elevated expression in the roots of genes involved in strigolactone biosynthesis (*NSP1*, *NSP2*, *CCD7*, and *MAX1a*), mycorrhiza-induced phosphate (*PT8*), ammonium (*AMT2;3*), and nitrate (*NPF4.12*) transporters and the putative ammonium transporter *NIP1;5*. *R. irregularis* also stimulated the expression of photosynthesis-related genes in the shoot and the upregulation of the sugar transporters *SWEET1.2, SWEET3.3* and *SWEET 12* and the lipid biosynthesis gene *RAM2* in the roots. In contrast, *G. aggregatum* induced the expression of biotic stress defense response genes in the shoots and several genes associated with abiotic stress in the roots. This suggests that either the host perceives colonization by *G. aggregatum* as a pathogen attack or that *G. aggregatum* can prime host defense responses. Our findings reveal novel insights into the molecular mechanisms that control the host plant response to colonization with high- and low-benefit arbuscular mycorrhizal fungal symbionts.

## INTRODUCTION

Nitrogen (N) and phosphorus (P) are the main nutrients that limit plant growth in natural and agricultural ecosystems (LeBauer and Treseder 2008; Vitousek et al. 2010). Conventional management practices typically address both N and P limitations through the application of chemical fertilizers, but fertilizer production, transportation, and application are both economically and environmentally expensive (Havlin et al. 2014). Implementing more sustainable methods to mitigate nutrient deficiencies in crops is essential to prevent continued environmental degradation and to still meet the nutritional demands of a growing human population.

To overcome nutrient limitations, plants have evolved the capacity to form alliances with mutualistic, root-associated microbes (Martin et al. 2017). The most widespread mutualism is the arbuscular mycorrhizal (AM) symbiosis formed by 72% of all known plant species and a relatively small number of fungal species in the Glomeromycotina (Brundrett and Tedersoo 2018; Spatafora et al. 2016). AM fungi provide the host plant with mineral nutrients, such as P and N, and improve the resistance of plants against abiotic and biotic stressors (Bücking and Kafle 2015; Kafle et al. 2019). Under P deficiency, plants increase the production of strigolactones, a class of phytohormones that also serve as signaling molecules perceived by AM fungi (Yoneyama et al. 2007). Strigolactones are exuded into the rhizosphere through the ATP-binding cassette transporter PDR1 (Kretzschmar et al. 2012). AM fungal perception of strigolactones triggers spore germination, hyphal branching, and the production of short-chain chitin oligomers (Besserer et al. 2008; Genre et al. 2013). These, along with lipo-chitooligosaccharides (Maillet et al. 2011; Rush et al. 2020), comprise the so-called that prime the host plant for colonization by activating the “common symbiosis signaling pathway" (CSSP; reviewed in MacLean et al., 2017).

The key steps of the CSSP include: 1) the perception of Myc factors at the plasma membrane of root epidermal cells by lysine-motif receptor-like kinases (*e.g.*, NFP and LYK3; Feng et al., 2019); 2) the activation of the secondary messenger mevalonate that likely activates multiple nuclear membrane-localized calcium channels (CASTOR, DMI1/POLLUX, and CNGC15); 3) oscillations in calcium ion concentration in the nucleoplasm that activate a calcium- and calmodulin-dependent protein kinase (DMI3/CCaMK; Lévy et al., 2004; Singh and Parniske, 2012); 4) phosphorylation of IPD3 which, in concert with DELLA, induces the expression of RAM1; and 5) activation of several other transcription factors required for the development and regulation of the AM symbiosis (Pimprikar et al., 2016; Pimprikar and Gutjahr, 2018). Following root penetration, AM hyphae proliferate both intra- and intercellularly and form highly branched nutrient exchange structures called arbuscules in cortical cells (Park et al. 2015). Surrounding the arbuscule is a plant-derived periarbuscular membrane (PAM) that is enriched in AM-induced transporters that facilitate reciprocal nutrient exchange between the fungus and the plant (Garcia et al. 2016). In *Medicago truncatula*, three AMT2 family ammonium transporters (AMT2;3, AMT2;4, and AMT2;5; Breuillin-Sessoms et al., 2015) and the phosphate transporters *PT4* (Javot et al. 2007) and *PT8* (Breuillin-Sessoms et al. 2015) are involved in the uptake of N and P from the periarbuscular space between the fungal plasma membrane and the PAM. In exchange, host plants transfer about 20% of their photosynthetically derived carbon to their fungal partners, and these high carbon costs force host plants to strongly regulate the carbon flux to their fungal symbionts (Wright et al. 1998). The host plant releases lipids and sugars into the mycorrhizal interface likely via the putative lipid transporters *STR* and *STR2* (Zhang et al., 2010; Roth and Paszkowski, 2017) and the bidirectional sugar transporter *SWEET1.2* (formerly SWEET1b, An et al., 2019; Doidy et al., 2019).

AM fungi differ in the benefits that they provide for their host plant. The cooperative AM fungus *Rhizophagus irregularis* 09 (Ri09) transfers more N and P to its host plant and contributes to higher mycorrhizal growth responses than the less-cooperative fungus *Glomus aggregatum* 165 (Ga165; Fellbaum et al., 2014). Symbiosis with Ga165 is also more costly for the host plant (measured as carbon costs per P transferred) than with Ri09, and *M. truncatula* preferentially allocates more carbon to Ri09 than to Ga165 (Kiers et al. 2011). Collectively, these studies suggest that carbon to nutrient exchange processes are driven by biological market dynamics, but the molecular mechanisms that regulate these processes are largely unknown. We hypothesized that the host plant regulates the expression of key genes in response to the colonization of AM fungi that differ in the nutrient benefits that they provide for the host. To test this hypothesis, we examined the physiological and transcriptomic responses of the host plant *M. trancatula* to colonization by either high- (Ri09) or low-benefit (Ga165) AM fungal partners under N and P limitation.

## RESULTS

### Colonization by *Rhizophagus irregularis* 09 leads to greater plant biomass and phosphorus and nitrogen uptake than *Glomus aggregatum* 165

We used a dual-compartment pot system to compare the overall benefit that Ga165 and Ri09 provided to *M. truncatula* (**Supplemental Fig. S1**). The percent root length colonization by intraradical hyphae and arbuscules was comparable in roots colonized by either fungus; however, the percent of roots containing vesicles was significantly higher in Ga165-colonized roots (**Fig. 1A**). Arbuscular width was comparable for both species of AM fungi (**Fig. 1B-D**). Compared to non-mycorrhizal (NM) plants, Ri09 colonization resulted in greater shoot and root biomass, whereas Ga165 provided no growth benefits (**Fig. 2A-B**). Both fungi led to increased P content (**Fig. 2C-D**) and concentration (**Supplemental Fig. S2**) in the shoots and roots of their host, but Ri09 transferred more P than Ga165. Nitrogen-15 (^15^N) enrichment levels in the roots and shoots of NM plants did not exceed natural abundance levels (**Fig. 2E-F**), which confirms that there was no mass flow of ^15^N from the HC into the RC. Both fungi transferred ^15^N from the HC to their host, but Ga165 only increased root ^15^N enrichment, whereas Ri09 increased both root and shoot ^15^N enrichment. Overall, these data indicate that although Ga165 and Ri09 showed similar levels of root colonization, only Ri09 colonization led to increased growth, most likely by providing the host plant with greater access to both P and N.

**Figure 1.**
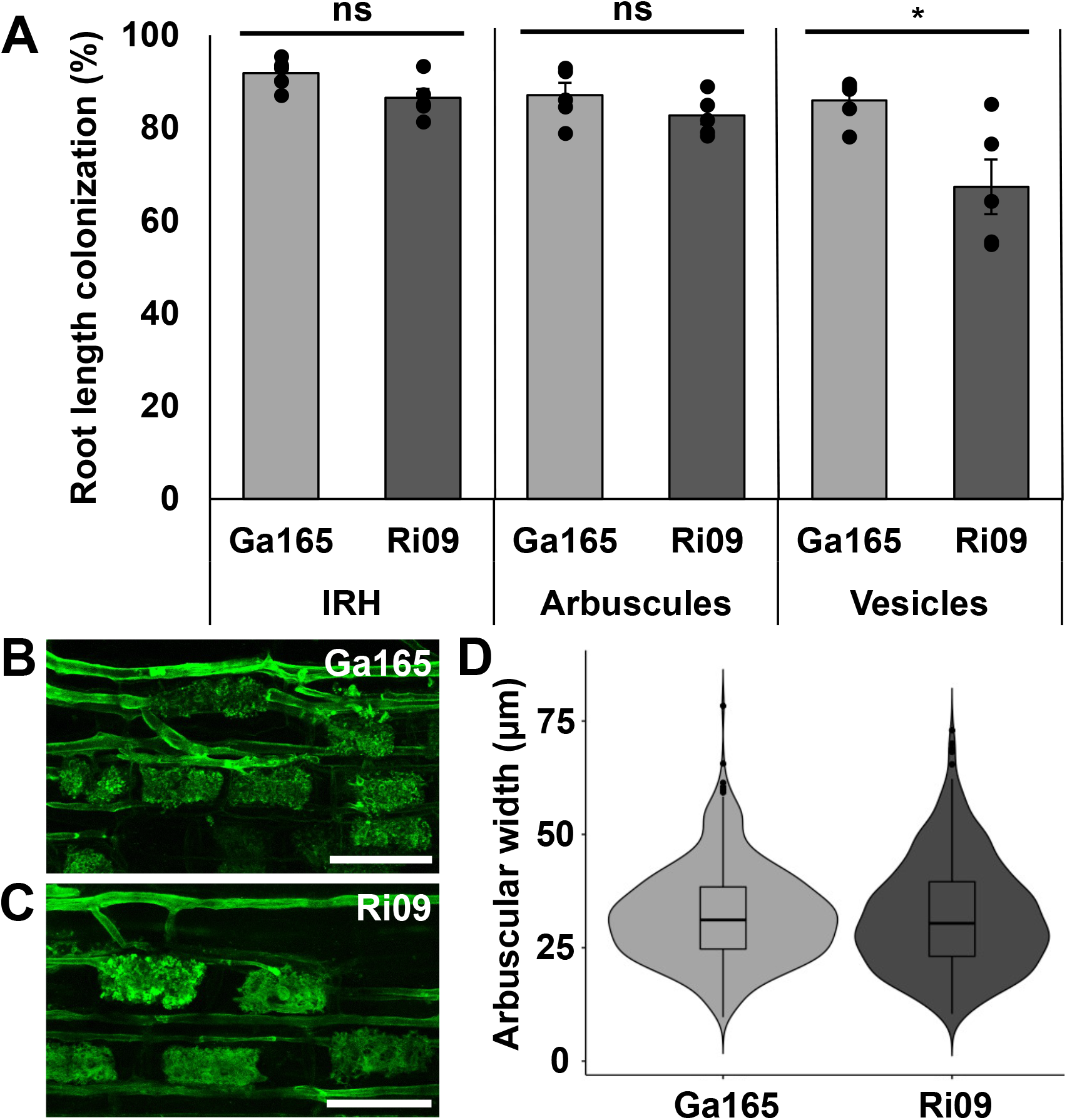
Arbuscular mycorrhizal colonization of *Medicago truncatula* roots and arbuscule size. **A,** percent root length colonization of roots inoculated with *Glomus aggregatum* 165 (Ga165; grey) or *Rhizophagus irregularis* 09 (Ri09; dark grey), measured for intraradical hyphae (IRH), arbuscules, and vesicles. Data points represent individual values for biological replicates (n = 5) and bars represent the mean ± SEM. **B-C**, representative confocal microscope images of Ga165 (**B**) and Ri09 (**C**) arbuscules stained with wheat germ agglutinin conjugated to Alexa Fluor 488. Scale bars = 50 μm. **D**, violin plot of arbuscule widths measured in roots colonized by Ga165 (grey) or Ri09 (dark grey) overlayed with box and whisker plots. In total, 488 and 508 arbuscules were measured in Ga165 and Ri09-colonized roots, respectively. All data were analyzed using a Welch’s twosample t-test and statistically significant differences (p ≤ 0.05) are indicated with an asterisk.

**Figure 2.**
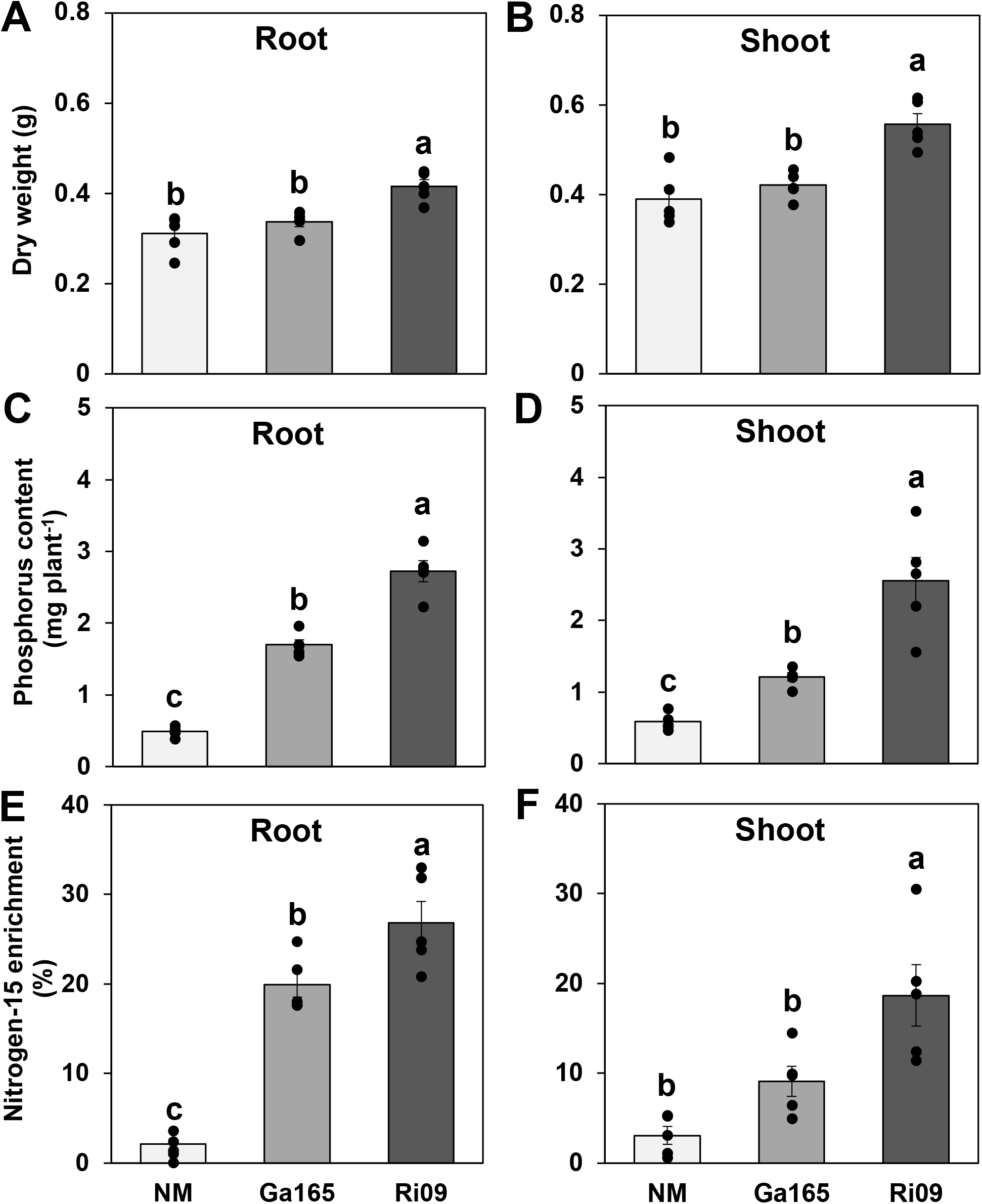
Biomass and nutrient content of *Medicago truncatula* plants colonized by different arbuscular mycorrhizal fungi. Shown are dry weight (**A**,**B**), phosphorus content (**C**,**D**), and % nitrogen-15 enrichment (**E**,**F**) of roots (left) and shoots (right). The plants were inoculated with either *Glomus aggregatum* 165 (Ga165) or *Rhizophagus irregularis* 09 (Ri09) and compared to non-mycorrhizal (NM) control plants. Data points represent individual values for biological replicates (n = 5) and bars represent the mean of each treatment ± SEM. Different letters on the bars indicate statistically significant (p ≤ 0.05) differences based on one-way ANOVA and LSD test. F statistics and p-values are shown in **Supplemental Table S1.**

### Colonization by either fungus alters gene expression more dramatically in the roots than in the shoots of *Medicago truncatula*

To identify potential molecular mechanisms responsible for the observed physiological responses, we used RNA sequencing (RNA-seq) to evaluate gene expression patterns in roots and shoots of mycorrhizal and non-mycorrhizal plants (**Supplemental File 1**). Analysis of the root data revealed distinct, treatment-specific hierarchical clustering among biological replicates (n=3, Ga165 and NM; n=4, Ri09) using a dendrogram, multidimensional scaling, and principal component analyses (**Supplemental Fig. S3**). In contrast, analysis of the shoot data showed no distinct clustering among biological replicates (n=4, Ga165 and NM; n=3, Ri09) within each treatment (**Supplemental Fig. S4**), suggesting that particularly in the roots, gene expression is driven by the fungal partner. Consequently, the overall number of significant differentially expressed genes (DEGs) was lower in the shoots than in the roots (**Supplemental Fig. S5**).

### Both shared and distinct sets of genes are differentially regulated in *Medicago truncatula* by colonization with either fungus

To identify key differences in the response of the host plant to both fungal species, we sorted all DEGs from each treatment and used an arbitrary threshold of log_2_(fold-change) > 2 and a q-value < 0.05 (**Fig. 3A**). In the roots, this threshold revealed that 364 genes were upregulated in roots colonized by either fungus, and 197 or 165 genes were upregulated by colonization with only Ga165 or Ri09, respectively (**Fig. 3B**). Ga165 and Ri09 colonization led to the downregulation of 41 shared genes, and 172 and 77 unique genes, respectively. The clustering of DEGs allowed us to identify specific groups of similarly expressed genes (**Fig. 3C**). Clusters A and B are comprised of genes that were strongly downregulated in mycorrhizal roots, cluster C1 contains genes that were upregulated in mycorrhizal roots, clusters C2 and D include genes primarily upregulated by Ga165 colonization, and cluster C3 contains genes exclusively upregulated by Ri09 colonization (**Supplemental File 2)**.

**Figure 3.**
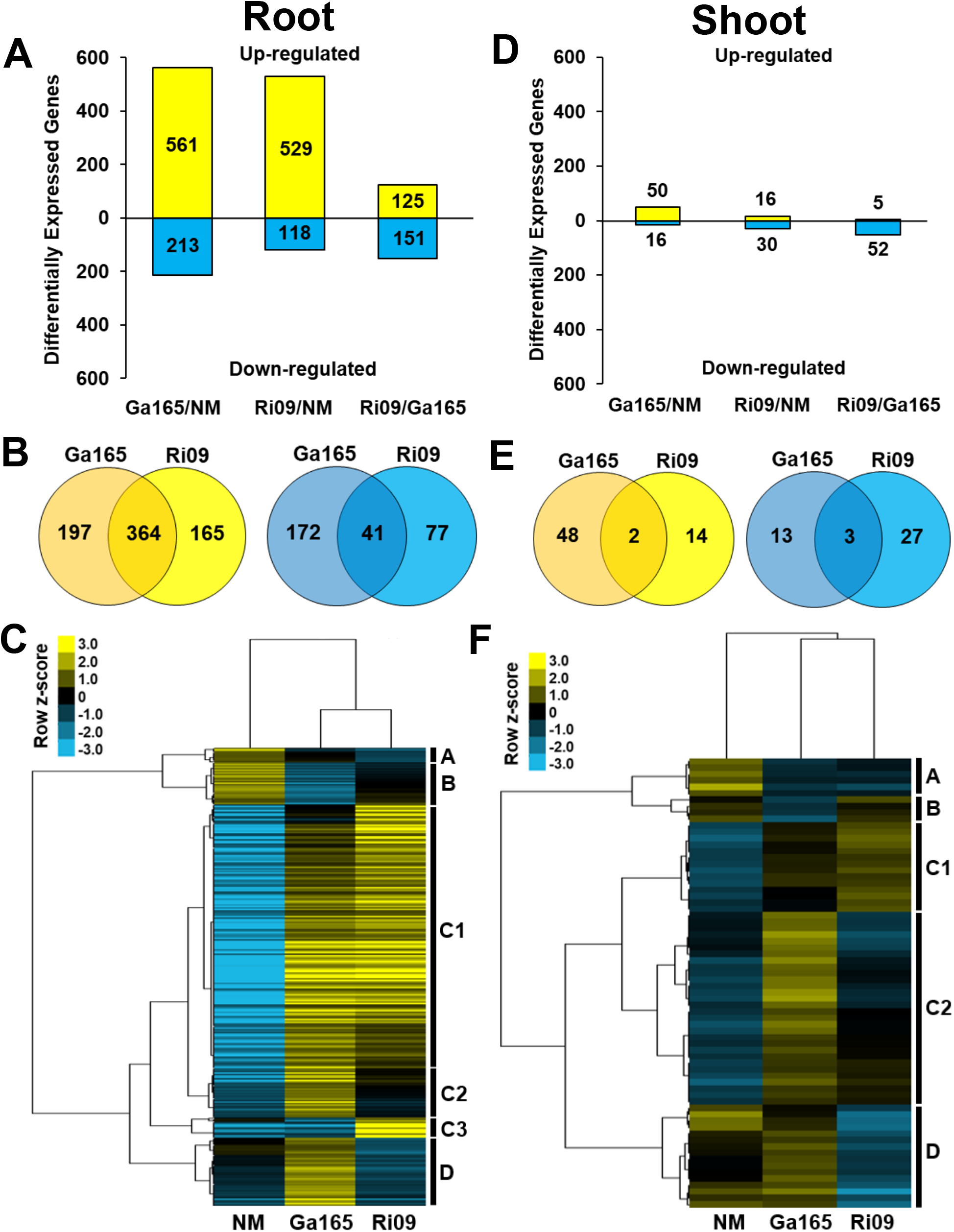
Significantly differentially expressed genes in roots (**A**–**C**) and shoots (**D**–**F**) of *Medicago truncatula* colonized by different AM fungi. **A** and **D**, Number of significantly differentially expressed genes (DEGs; log_2_[fold-change] > 2 and a q-value < 0.05) in plants colonized by *Glomus aggregatum* 165 (Ga165) or *Rhizophagus irregularis* 09 (Ri09) compared either to non-mycorrhizal (NM) control plants or to each other. **B** and **E**, Venn diagrams of significantly upregulated (yellow) and downregulated (blue) DEGs unique to Ga165 and Ri09-colonized plants or shared by both. **C** and **F**, Heat maps of fragments per kilobase of transcript per million mapped reads (FPKM) values + 1 for all significant DEGs from each treatment. Gene clusters are assigned based on secondorder and sub-clusters based on third- or fourth-order branching. Upregulated genes are shown in yellow and downregulated genes in blue with intensity based on row z-score. All DEGs in each plot for roots and shoots, respectively, are listed in **Supplemental Files 2 and 3**.

Although fewer genes were differentially expressed in the shoots, many were above our applied threshold (**Fig. 3D**). We found that 48 and 14 genes were up, and 13 and 27 genes were downregulated in the shoots of Ga165 and Ri09-colonized plants, respectively (**Fig. 3E**). In contrast to the roots, clustering of DEGs based on expression patterns in the shoots (**Fig. 3F)** revealed that only a small number of genes were upregulated (cluster C1) or downregulated (cluster A) in mycorrhizal plants. In addition, some genes were only downregulated by Ga165 (cluster B) or by Ri09 colonization (cluster D), while others were primarily upregulated by Ga165 colonization (cluster C2; **Supplemental File 3**). Collectively, these results demonstrate that compared to shoots, gene expression patterns in roots respond more strongly to colonization with different AM fungi, and that both fungi lead to distinct differences particularly in the root and to a lesser degree also in the shoot transcriptome profiles.

### Genes critical for the establishment of AM symbiosis are more strongly upregulated by *Rhizophagus irregularis* 09 than *Glomus aggregatum* 165

We conducted a targeted analysis of the root RNA-seq data by first evaluating the expression of many genes known to play a role in AM symbiosis (**Supplemental Table S2**). Among six core strigolactone biosynthesis or exporter genes (**Fig. 4A**), we observed that two (*DXS2* and *D27*) were upregulated in AM roots compared to the NM control; one (*MAX1a*) was only upregulated in Ri09-colonized roots, while three (*CCD7*, *CCD8-1* and *PDR1a*) were not affected. Two GRAS transcription factors (*NSP1* and *NSP2*) that regulate strigolactone biosynthesis and other symbiosis genes were also upregulated, particularly in Ri09-colonized roots. The combined increased expression of *NSP1*, *NSP2*, *CCD7*, and *MAX1a* in Ri09-colonized roots compared to Ga165-colonized roots suggests that Ri09 more strongly induces strigolactone biosynthesis than Ga165.

**Figure 4.**
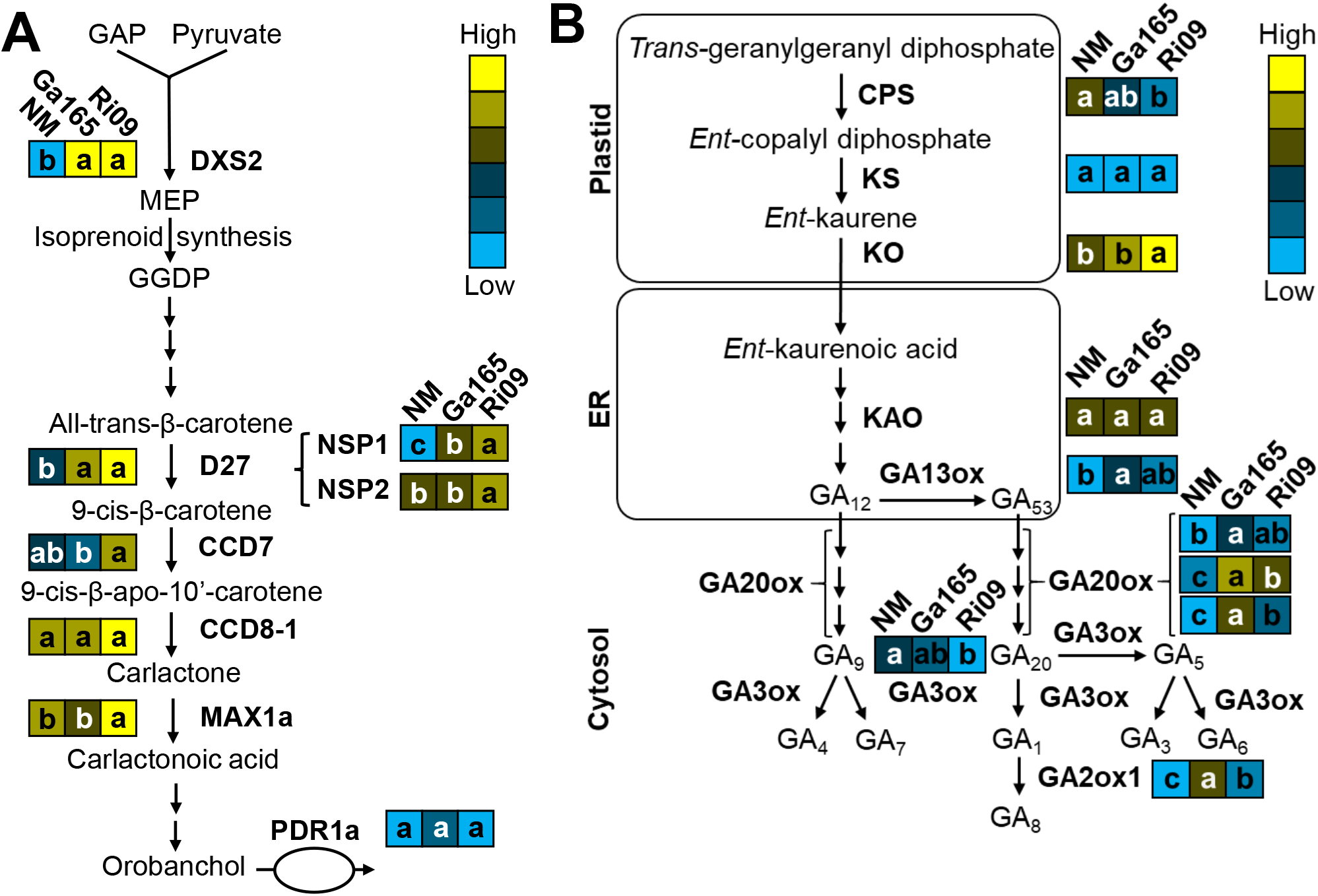
Expression patterns of genes encoding enzymes involved in the strigolactone (**A**) and gibberellic acid (**B**) biosynthesis pathways in the roots of *Medicago truncatula* colonized by arbuscular mycorrhizal fungi. Each diagram portrays the relative expression levels of genes that encode components of the pathways shown as fragments per kilobase of transcript per million mapped reads (FPKM; blue=low, yellow=high) in roots from non-mycorrhizal (NM) plants or plants colonized by either *Glomus aggregatum* 165 (Ga165) or *Rhizophagus irregularis* 09 (Ri09). Letters within boxes indicate significant differences (q-value < 0.05) in FPKM values between treatments and were determined as part of the CuffDiff2 analysis (see **Supplemental Table S2**). Abbreviations: **A**, GAP, glyceraldehyde 3-phosphate; DXS2, 1-deoxyxylulose 5-phosphate synthase2; MEP, 2-methyl-erythritol-4-phosphate; GGDP geranylgeranyl pyrophosphate; D27, Dwarf27; CCD, carotenoid cleavage dioxygenase; and PDR1a, pleiotropic drug resistance 1a. **B**, CPS, *ent*-copalyl diphosphate synthase; KS, *ent*-kaurene synthase; KO, *ent*-kaurene oxidase; KAO, *ent*-kaurenoic acid oxidase; GA13ox, gibberellin 13-oxidase; GA20ox, gibberellin 20-oxidase; GA3ox, gibberellin 3-oxidase; and GA2ox, gibberellin 2-oxidase.

Among the CSSP genes, the lipochitooligosaccharide receptor *NFP* was downregulated and *PUB1*, a negative regulator of AM symbiosis (Vernié et al. 2016), was upregulated in roots colonized by either fungus (**Supplemental Table S2**). None of the other 13 analyzed CSSP genes were affected. However, gibberellic acid (GA) is known to play an important role in regulating AM colonization (Floss et al. 2013; Nouri et al. 2021), and several genes regulating the biosynthesis and degradation of GA were differentially expressed by mycorrhizal colonization (**Fig. 4B**). In Ri09-colonized roots, the expression of the genes encoding *ent*-copalyl diphosphate synthase (*CPS*) and GA3oxidase (*GA3ox*) were down-regulated compared to NM roots, but the gene encoding *ent*-kaurene oxidase (*KO*) was upregulated. In Ga165-colonized roots, four GA biosynthesis genes (*GA13ox* and three homologs of *GA20ox*) were significantly upregulated compared to NM roots. In Ri09-colonized roots also two of the *GA20ox* homologs were upregulated, but not as strongly as by GA165. Similarly, the GA degradation gene *GA2ox1* was also upregulated in mycorrhizal roots, but more strongly in Ga165-colonized roots (**Supplemental Table S2**).

Given the higher expression of more GA biosynthesis genes in Ga165 compared to Ri09-colonized roots, we hypothesize that GA plays a stronger role in regulating the symbiosis with GA165. Therefore, we evaluated the expression of genes downstream of of the CSSP and DELLA1/2, which are proteins that are negatively regulated by GA (Davière and Achard 2013). We evaluated the expression of 21 genes (**Supplemental Table S3**), including: seven AM-induced transcription factors that regulate arbuscule development, function, or degradation — *DIP1*, *RAD1*, *RAM1*, *MIG1*/*2*/*3*, and *MYB1*; three cellular remodeling genes — *EXO70I*, *SYP132A*, and *VAPYRIN*; two phosphate transporters (*PT4* and *PT8*); three ammonium transporters (*AMT2;3*, *AMT2;4*, and *AMT2;5*); the putative lipid transporters *STR* and *STR2* and three lipid biosynthesis genes — *RAM2*, *FatM*, and *WRI5a*; and the sugar transporter *SWEET1.2*. Almost all these genes were upregulated by both AM fungal species; however, six were substantially more upregulated by Ri09, including: *RAD1*, *MYB1*, *PT8*, *AMT2;3*, *RAM2*, and *SWEET1.2* (**Fig. 5A-F**). Together with the physiological results, the increased upregulation of *PT8*, *AMT2;3*, and *SWEET1.2* in Ri09-colonized roots is suggestive of greater nutrient exchange activity across the PAM than in Ga165-colonized roots.

**Figure 5.**
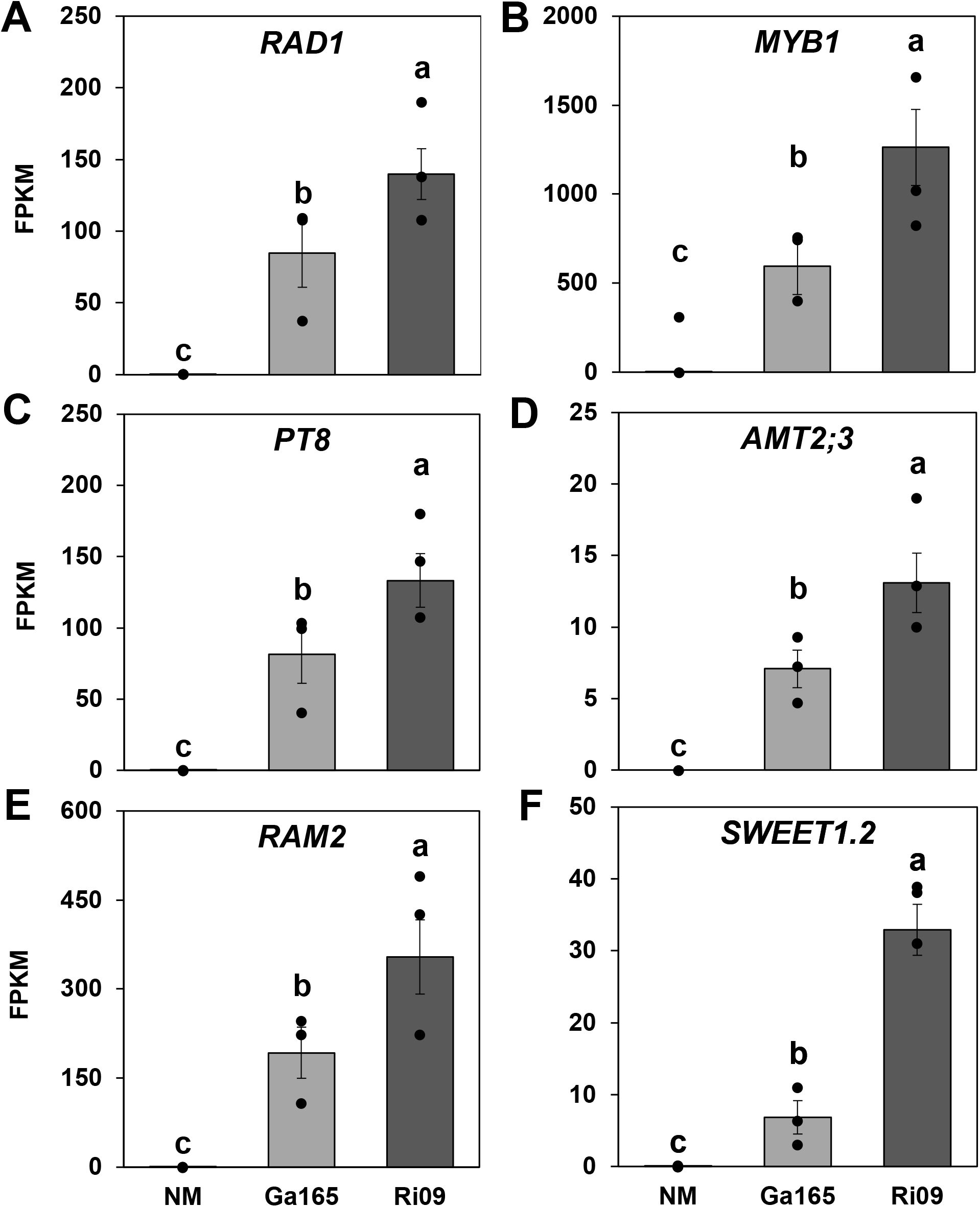
Relative expression of six genes in the roots of *Medicago truncatula* after colonization with different arbuscular mycorrhizal fungi. Expression of **A**, *Required for Arbuscule Development 1* (*RAD1*); **B**, *MYB-like Transcription Factor 1* (*MYB1*); **C**, *Phosphate Transporter 8* (*PT8*); **D**, *Ammonium Transporter 2;3* (*AMT2;3*); **E**, *Reduced Arbuscular Mycorrhization 2* (*RAM2*); and **F**, *Sugars Will Eventually Be Exported Transporter 1.2* (*SWEET1.2*) in the roots of non-mycorrhizal plants (NM, n=3) and plants inoculated with *Glomus aggregatum* 165 (Ga165, n=3) or *Rhizophagus irregularis* 09 (Ri09, n=4). Expression levels are shown as fragments per kilobase of transcript per million mapped reads (FPKM). Data points represent individual values for biological replicates (n = 5) and bars represent the mean of each treatment ± SEM. Significant differences (qvalue < 0.05) in FPKM values between treatments were determined as part of the CuffDiff2 analysis and are indicated using different letters (see **Supplemental Table S3**).

### Colonization with *Glomus aggregatum* 165 and *Rhizophagus irregularis* 09 differentially affect the expression of transporters

Gene ontology (GO) enrichment analysis revealed several cellular components, molecular functions, and/or biological processes that were significantly enriched among genes that are only upregulated (**Supplemental Fig. S6**) or downregulated in AM roots (**Supplemental Fig. S7**). Predominant among these were GO terms related to transmembrane transport. We from the *M. truncatula* JCVI 4.0v2 annotation file (Tang et al. 2014) and found 259 genes that were significantly differentially regulated in mycorrhizal roots (**Supplemental File 5**). Based on their function, we further subdivided these genes into two groups: transport of mineral nutrients (72 genes) and secondary metabolites (187 genes). Compared to NM roots, 12 mineral transporters were up- and 17 were down-regulated by both AM fungi (**Fig. 6A**). This suggests that a core set of mineral transporters in AM roots are similarly regulated independent of fungal benefits. However, a unique set of transporters were differentially regulated by each AM fungal species: 14 were up- and 13 were down-regulated in Ga165-colonized roots, while only 7 were up- and 12 were down-regulated in Ri09-colonized roots. From the identified 16 groups of mineral transporters (**Fig. 6B**), we focused on ammonium, nitrate, phosphate, and sulfate transporters with FPKM 10, due to the key role that AM fungi play in the transport of these nutrients (**Fig. 6C-F**). Among the five differentially regulated ammonium transporters, one belongs to the AMT1 ammonium transporter family and four to the AMT2 family. Compared to AM roots, *AMT1;1* and two of the AMT2 family transporters (*AMT2;1* and *AMT2;6*) were highly expressed in NM roots, while *AMT2;3* and *AMT2;5* were specifically induced in mycorrhizal roots. *AMT2;3* was particularly up-regulated in Ri09-colonized roots (**Fig. 6C**).

**Figure 6.**
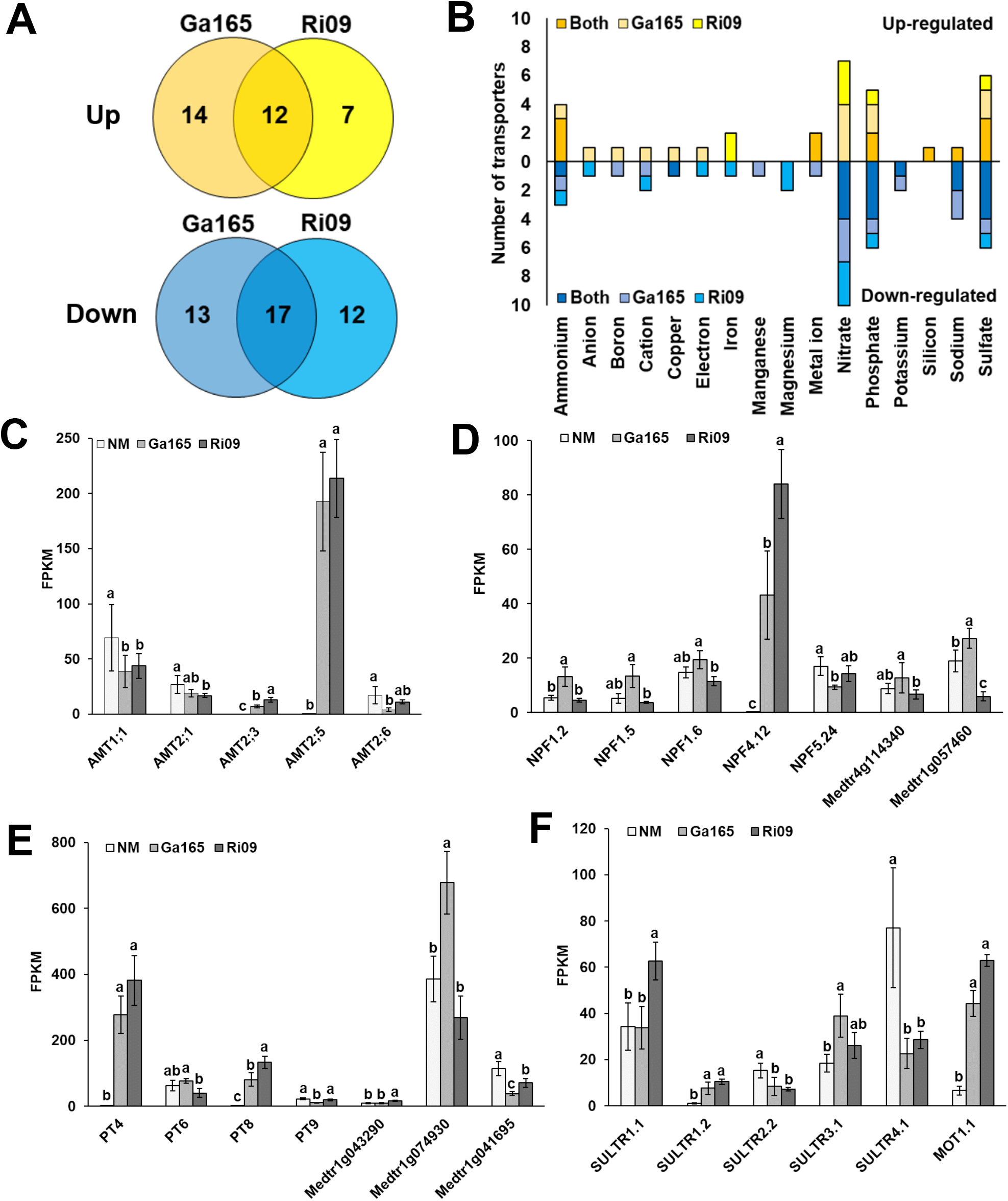
Expression patterns of significantly differentially expressed mineral transporter genes in the roots of *Medicago truncatula* after colonization with different arbuscular mycorrhizal fungi. **A**, Venn diagrams showing the number of mineral transporters that are significantly up-(yellow) or down-regulated (blue) in roots colonized by *Glomus aggregatum* 165 (Ga165) or *Rhizophagus irregularis* 09 (Ri09), or both. **B**, summary of the number of genes from specific gene annotation groups of mineral transporters that are up- or down-regulated in roots colonized by Ga165 or Ri09, or both (see also **Supplemental File 6**). Expression patterns of genes annotated as **C**, ammonium; **D**, nitrate; **E**, phosphate; and **F**, sulfate transporters in the roots of non-mycorrhizal plants (NM; n=3) and plants inoculated with Ga165 (n=3) or Ri09 (n=4). The bars in C–F represent the mean of each treatment ± SEM (data points for biological replicates are displayed in **Supplemental Figure S8**). Expression levels are shown as fragments per kilobase of transcript per million mapped reads (FPKM), only genes with FPKM ≥ 10 are shown, significant differences (q-value < 0.05) in FPKM values between treatments were determined as part of the CuffDiff2 analysis and are indicated using different letters on the bars.

We also identified seven differentially regulated nitrate transporters (**Fig. 6D**). Five belong to the nitrate/peptide transporter (NPF) family, and two are putative nitrate transporters. Interestingly, the most highly expressed nitrate transporter (*NPF4.12*) was highly upregulated in AM roots, particularly in Ri09-colonized roots. In contrast, five nitrate transporters were more highly expressed in Ga165-colonized roots than in Ri09-colonized roots (*NPF1.2*, *NPF1.5*, *NPF1.6*, *Medtr4g114340*, and *Medtr1g057460*), and two (*NPF1.2* and *Medtr1g057460*) also differed from control roots. Only the nitrate transporter *NPF5.24* and *Medtr1g057460* showed higher expression in NM roots than in AM-colonized roots.

Seven phosphate transporters were differentially regulated (**Fig. 6E**). Four belong to the high-affinity inorganic phosphate transporter family PHT1 (*PT4*, *PT6*, *PT8*, and *PT9*), two were annotated as high-affinity inorganic phosphate transporters (*Medtr1g043290* and *Medtr1g074930*), and one as a phosphate transporter PHO1-like protein. As expected, two of the PHT1 transporter genes, *PT4* and *PT8*, were only expressed in AM roots, and *PT8* expression was higher in Ri09-colonized roots. By contrast, the expression of *Medtr1g074930* was strongly upregulated in Ga165-colonized roots compared to NM and Ri09-colonized roots. The PHO1-like transporter (*Medtr1g041695*) showed higher expression levels in NM than in AM roots.

We identified six sulfate transporters that were differentially regulated among the three treatments (**Fig. 6F**). Two were annotated as high affinity sulfate transporter 1 (SULTR1) family genes (*SULTR1.1* and *1.2*), three as sulfate/bicarbonate/oxalate exchanger and transporter genes (*SULTR2.2*, *3.1*, and *4.1*), and one as a sulfate transporter-like gene (*MOT1.1*). While the expression levels of *SULTR2.2* and *SULTR4.1* were higher in NM roots than in AM roots, the remaining sulfate transporters were upregulated by colonization with Ri09 (*SULTR1.1*), Ga165 (*SULTR3.1*), or by both fungi (*SULTR1.2* and *MOT1.1*).

In total, we identified 20 groups of secondary metabolite transporters in our dataset. Compared to NM control roots, 29 of these transporters were up- and 43 were downregulated in AM roots (**Fig. 7A**). Several secondary metabolite transporters were differentially regulated in Ga165 and Ri09-colonized roots. Among these, most belonged to six gene annotation groups, including ATP-binding cassette (ABC), amino acid, major facilitator superfamily (MFS), major intrinsic protein (MIP), peptide, and Sugars Will Eventually be Exported Transporters (SWEETs) (**Fig. 7B**). Here, we will only discuss those transporters that are known to play a role in AM symbiosis, or that were strongly differentially regulated by mycorrhizal colonization, but have not yet been functionally characterized.

**Figure 7.**
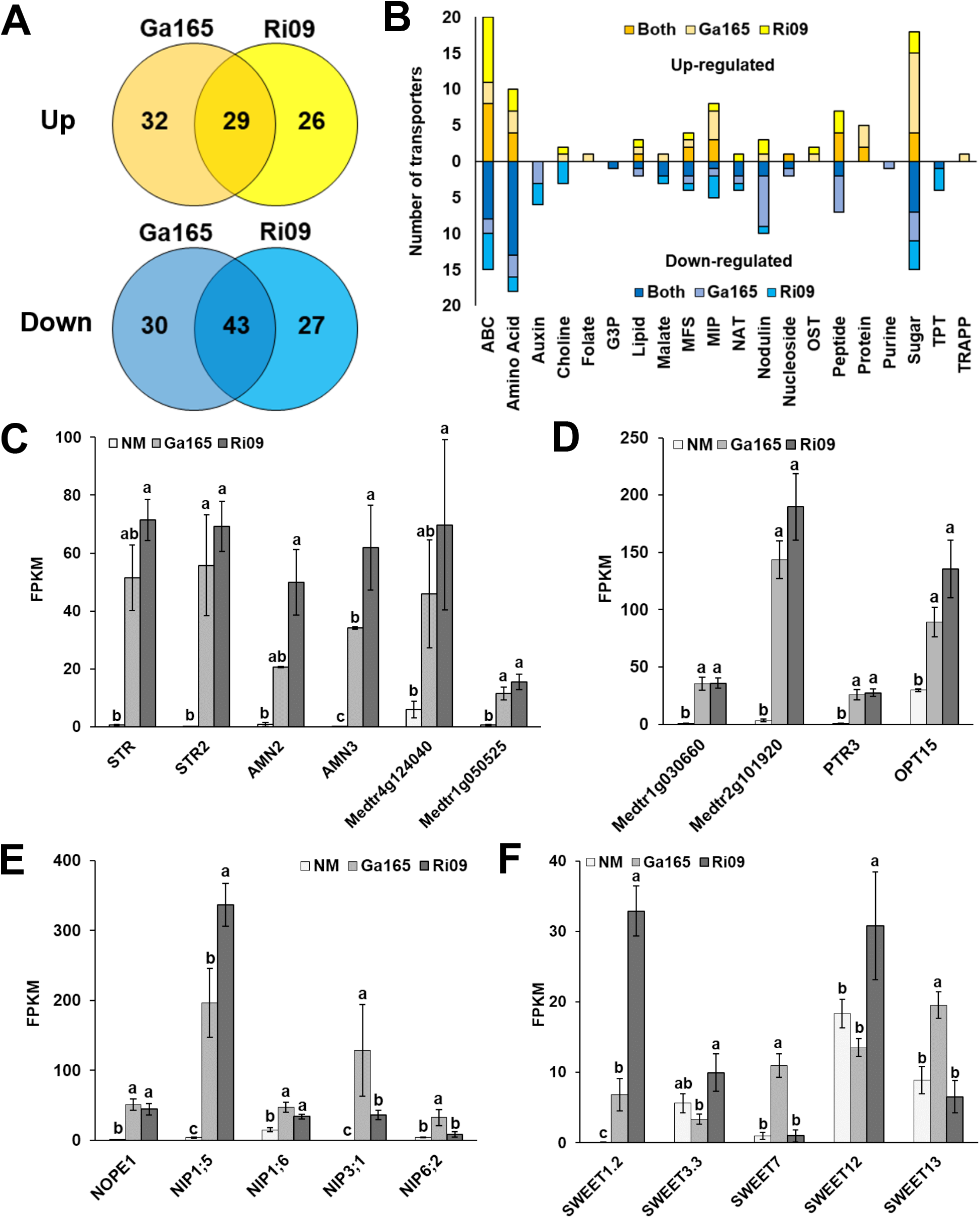
Expression patterns of significantly differentially expressed secondary metabolite transporters in the roots of *Medicago truncatula* after colonization with different AM fungi. **A**, Venn diagrams showing the number of secondary metabolite transporters that are significantly up- (yellow) or down-regulated (blue) in roots colonized by *Glomus aggregatum* 165 (Ga165) or *Rhizophagus irregularis* 09 (Ri09), or both. **B**, summary of the number of genes from specific gene annotation groups of secondary metabolite transports that are up- or down-regulated in roots colonized by Ga165 or Ri09, or both (see **Supplemental File 7**). Expression patterns of genes annotated as **C**, ATP-binding cassette (ABC); **D**, amino acid or peptide; **E**, major facilitator superfamily (MFS) or major intrinsic protein (MIP); and **F**, SWEET transporters in the roots of non-mycorrhizal plants (NM; n = 3) and plants inoculated with Ga165 (n =3) or Ri09 (n = 4). The bars in C-F represent the mean of each treatment ± SEM (data points for biological replicates are displayed in **Supplemental Figure S9**). Expression levels are shown as fragments per kilobase of transcript per million mapped reads (FPKM); only genes with FPKM ≥ 10 are shown; significant differences (q-value < 0.05) in FPKM values between treatments were determined as part of the CuffDiff2 analysis and are indicated using different letters on the bars. Abbreviations in B: G3P, glycerol-3-phosphate; NAT, nucleobase-ascorbate transporter; OST, organic solute transporter; TPT, triose-phosphate transporter; TRAPP, transport protein particle.

Two white-brown-complex ABC transporter family proteins (*STR* and *STR2*; Zhang et al., 2010), three ABC transporter B family proteins (*AMN2*, *AMN3* [Roy, 2015], and *Medtr4g124040*), and one drug resistance transporter-like ABC domain protein (*Medtr1g050525*) were strongly upregulated in AM roots, particularly in Ri09-colonized roots (**Fig. 7C**). The expression levels in Ga165-colonized roots were higher but did not always differ significantly from the NM roots.

Two amino acid transporters, *Medtr1g030660* and *Medtr2g101920*, and two peptide transporters, *PTR3* and *OPT15*, were upregulated in mycorrhizal roots (**Fig. 7D**). As expected, *NOPE1,* an N-acetylglucosamine transporter of the major facilitator superfamily (MFS) that is required for AM symbiosis (Nadal et al. 2017), was also highly upregulated in AM roots (**Fig. 7E**). Three major intrinsic protein (MIP) transporters, *NIP1;5*, *NIP1;6*, and *NIP3;1*, which are often classified as aquaporins, were all upregulated in mycorrhizal roots. *NIP1;5* was particularly highly expressed in Ri09-colonized roots, while *NIP1;6*, *NIP3;1* and *NIP6;2* showed higher expression levels in Ga165-colonized roots.

SWEETs are known to play a crucial role in AM symbiosis (Doidy et al. 2019), and five were differentially expressed in AM roots. *SWEET1.2* was upregulated by colonization with either fungus, but especially with Ri09. *SWEET3.3* and *SWEET12,* were only upregulated in Ri09-colonized roots, whereas *SWEET7* and *SWEET13* were only upregulated in Ga165-colonized roots (**Fig. 7F**). The variation in the expression patterns of these five *SWEET* genes suggests that sugar transport is strongly regulated in response to fungal benefit.

### Photosynthesis-related genes in the shoots are upregulated in response to colonization with *Rhizophagus irregularis* 09

GO enrichment analysis on several gene clusters shown in **Fig. 2F** revealed that cellular components and biological processes associated with photosynthesis were enriched in the shoots of Ri09-colonized plants (**Supplemental Fig. S10A**). The associated genes were annotated as follows: *Medtr1g115410*, photosystem II reaction center family protein (orthologous to PsbP in *Arabidopsis thaliana*); *Medtr2g082580*, oxygen-evolving enhancer protein (orthologous to PsbQ-like protein 2 in *A. thaliana*); *Medtr5g018670*, photosystem II oxygen-evolving enhancer protein; *Medtr1g015290*, ultraviolet-B-repressible protein (orthologous to PsbX in *A. thaliana*); and *Medtr1588s0010*, ATP synthase F1, gamma subunit. The expression levels of *PsbP*, *PsbQ-like protein 2*, *PsbX*, and the ATP synthase gene were significantly higher in the shoots of Ri09-colonized plants than in NM plants (**Fig. 8A**). Out of these genes, only *PsbP* was also significantly upregulated in Ga165-colonized plants. By contrast, *Medtr5g018670* was more highly expressed in the shoots of Ga165-colonized than in NM plants. Overall, these expression patterns imply that photosynthesis was more strongly induced in the shoots of Ri09-colonized plants, which could partially explain the increased biomass of Ri09-colonized plants.

**Figure 8.**
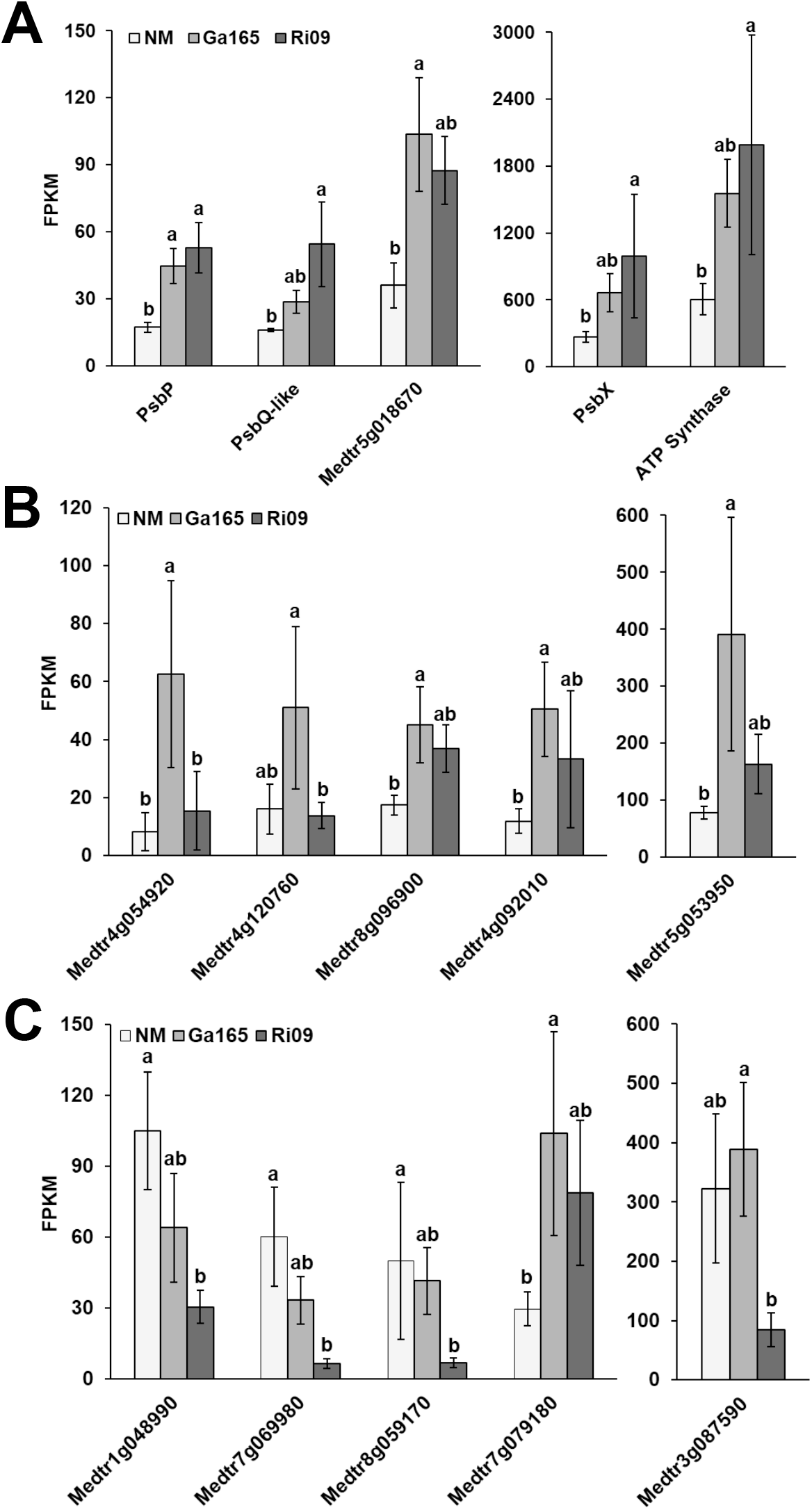
Expression patterns of significantly differentially expressed genes in the shoots of *Medicago truncatula* colonized by different AM fungi. Expression levels of genes involved in **A**, photosynthesis; **B**, response to biotic stress; and **C**, response to abiotic stress in non-mycorrhizal plants (NM; n=4) or plants colonized by *Glomus aggregatum* 165 (Ga165, n=4) or *Rhizophagus irregularis* 09 (Ri09, n=3). Bars represent the mean of each treatment ± SEM (data points for biological replicates are displayed in **Supplemental Figure S11**). Expression levels are shown as fragments per kilobase of transcript per million mapped reads (FPKM), and significant differences (q-value < 0.05) in FPKM values between treatments were determined as part of the CuffDiff2 analysis and are indicated using different letters above the bars (see also **Supplemental Table S4**).

### Biotic and abiotic stress-related genes are differentially regulated in the shoots of *Glomus aggregatum* and *Rhizophagus irregularis* colonized plants

Many of the GO terms that were enriched in the shoots of Ga165-colonized plants encompassed biological processes related to biotic stress (**Supplemental Fig. S10B**). The genes linked to these GO terms were annotated as: *Medtr4g054920*, cytochrome P450 family 94 protein; *Medtr4g120760*, pathogenesis-related protein bet V I family protein; *Medtr8g096900,* pathogenesis-related thaumatin family protein; *Medtr4g092010*, (3S)-linalool/(E)-nerolidol/(E,E)-geranyl linalool synthase; and *Medtr5g053950*, allene oxide cyclase. Enrichment of these GO terms in the shoots of Ga165-colonized plants could indicate that the host plant is responding to Ga165 as it would to a pathogen by upregulating defense responses, or that Ga165 stimulates host plant defenses against pathogens (**Fig. 8B**).

In contrast, Ri09 colonization led to the downregulation of genes (**Fig. 3F**, cluster D), associated with biotic and abiotic stress (**Supplemental Fig. S10C**). The genes included in these GO terms were annotated as: *Medtr1g048990*, superoxide dismutase; *Medtr7g069980*, ferritin; *Medtr8g059170*, NAC transcription factor-like protein; *Medtr7g079180*, late embryogenesis abundant protein; and *Medtr3g087590*, myo-inositol 1-phosphate synthase. The expression of *Medtr7g079180* was upregulated by Ga165, but the remaining abiotic stress response genes were more highly expressed in NM or Ga165-colonized plants than in Ri09-colonized plants (**Fig. 8C**). Collectively, the expression patterns of these abiotic stress response genes could indicate that Ri09 is more effective at conferring abiotic stress tolerance to *M. truncatula* than Ga165.

## DISCUSSION

### The growth and nutritional benefits of arbuscular mycorrhizal fungi are linked to mycorrhiza-induced transporter expression

When compatible host plants are inoculated with AM fungi, colonization most often leads to increased growth and nutrient uptake, especially in nutrient-deprived conditions (Chandrasekaran 2020). We found that under N and P limitation, Ga165 and Ri09 colonized *M. truncatula* equally well (**Fig. 1**), but still, Ri09 promoted greater growth and provided more N and P to the host than Ga165 (**Fig. 2**). This observation is consistent with both the expression patterns that we observed for mycorrhiza-induced N and P transporters and with previous findings where we demonstrated that Ri09 is more cooperative than Ga165 (Kiers et al. 2011; Fellbaum et al. 2014; Wang et al. 2016). For example, the known AM-induced ammonium transporters *AMT2;3* and *AMT2;5* were both upregulated by Ga165 and Ri09 (**Fig. 6C**). Although the expression levels of *AMT2;3* in AM roots were lower than that of *AMT2;5*, *AMT2;3* expression was more strongly upregulated by Ri09 than by Ga165 colonization. Similarly, we observed the same expression pattern for the putative ammonium transporter *NIP1;5* (**Fig. 7E**), which has not been functionally characterized in *M. truncatula* during AM symbiosis, but the soybean ortholog *Nod26* plays a crucial role in ammonium transport in the rhizobia-legume symbiosis (Frare et al. 2018). Interestingly, although the AM fungi were provided with ^15^NH_4_Cl as a N source, we also found that the nitrate transporter *NPF4.12* was strongly induced in AM roots, especially in Ri09-colonized roots (**Fig. 6D**). Aloui et al. (2018) found that *NPF4.12* is enriched in the proteome of mycorrhizal *M. truncatula* roots, and recently the rice ortholog *OsNPF4.5* was functionally characterized as an AM-induced nitrate transporter (Wang et al. 2020). Overall, the stronger expression of ammonium and nitrate transporters in Ri09-than in Ga165-colonized roots aligns with the observed increase in the ^15^N labeling in the shoots of Ri09-colonized plants.

The AM-induced P transporters *PT4* and *PT8* were both highly expressed in AM colonized roots, especially Ri09-colonized roots (**Fig. 6E**). Similarly, colonization with both fungi, but particularly with Ri09, caused an increase in shoot P content. However, *Medtr1g074930*, a putative high affinity inorganic phosphate transporter, was strongly upregulated in Ga165-colonized roots (**Fig. 6E**). This P transporter is upregulated during P deficiency (Wang et al. 2017), and the upregulation of *Medtr1g074930* in our study could indicate that colonization by Ga165 resulted in P deficiency in the host plant. However, this is inconsistent with the observation that *Medtr1g074930* was not upregulated in the roots of NM plants (**Fig. 6E**) and Ga165-colonized roots had a higher P concentration than NM roots (**Supplemental Fig. S2**). Regardless, others have similarly observed that the expression of high affinity P transporters are differentially regulated in the host by colonization with different species of AM fungi (Grunwald et al. 2009).

Like N and P, AM fungi also play a critical role in supplying sulfur to the host plant (Sieh et al. 2013). We observed an upregulation of two sulfate transporters, *SULTR1.1* and *SULTR1.2* (**Fig. 6F**). Both are upregulated in mycorrhizal *M. truncatula* roots, although not consistently (Casieri et al. 2012; Wipf et al. 2014). The *SULTR1.2* ortholog in *Lotus japonicus* appears to play an important role in both direct and mycorrhizal sulfur uptake pathways, and its promoter is active in arbuscule-containing cells (Giovannetti et al. 2014). We also observed that the putative sulfate transporter gene *MOT1.1* was upregulated in Ga165 and Ri09-colonized roots (**Fig. 6F**). However, the involvement of MOT1.1 in sulfur transport has not been experimentally demonstrated yet, but it is phylogenetically related to *MOT1.3*, which is a molybdate transporter crucial for the rhizobia-legume symbiosis (Tejada-Jiménez et al. 2017).

### Patterns of photosynthesis, carbon transporter, and lipid biosynthesis gene expression in the host plant are linked to the nutritional benefits provided by the fungal symbiont

In exchange for the nutritional benefits that AM fungi provide, plants transfer between 4 and 20% of their net fixed carbon in the form of carbohydrates and lipids to these fungal symbionts (Wright et al. 1998; Roth and Paszkowski 2017). Both the nutritional benefits that AM fungi provide and the increased carbon sink they create are known to stimulate an increase in photosynthesis (Kaschuk et al. 2009). In line with this, we observed that the expression levels of photosynthesis-related genes were more strongly induced in Ri09-than in Ga165-colonized plants (**Fig. 8A**); thus, the elevated growth response provided by Ri09 (**Fig. 2A-B**) was likely not only due to an increased nutrient transfer, but potentially also due to a stimulation in photosynthesis.

Our previous studies have shown that host plants preferentially allocate more carbon to the more cooperative AM fungus Ri09 (Kiers et al. 2011). Yet, we found that the putative lipid transporters *STR* and *STR2* were upregulated in AM roots, but did not differ significantly between Ga165 and Ri09-colonized roots (**Fig. 7C**). Nevertheless, we did observe that the expression levels of the lipid biosynthesis gene *RAM2* (**Fig. 5**) and the sugar transporter *SWEET1.2* (**Fig. 5F**) were higher in Ri09-colonized roots, which is suggestive of increased transport of carbon resources to the fungus in exchange for elevated nutritional benefits (Bravo et al. 2017; An et al. 2019).

Although *SWEET1.2* is highly upregulated during AM colonization, particularly in arbusculated cortical cells, it is not essential for the maintenance of the AM symbiosis, most likely because of functional redundancy between SWEET1.2 and other SWEETs (An et al. 2019). We found that *SWEET3.3* and *SWEET12* were more strongly upregulated in Ri09-colonized roots (**Fig. 7F**); both are upregulated in arbusculated cortical cells in *M. truncatula* (Sameeullah et al. 2016). Interestingly, *SWEET7* and *SWEET13* were more strongly induced in Ga165-colonized roots, but SWEETs perform diverse physiological functions, and their upregulation does not necessarily increase sugar transport to the fungus. For example, some SWEET transporters in *Arabidopsis* localize to the tonoplast and restrict sugar efflux from the root (Chen et al. 2015). Given the variable expression patterns of these five SWEETs in Ri09 and Ga165-colonized roots and the varying roles they may play in sugar efflux and retention, functionally characterizing them will improve our understanding of the mechanisms controlling carbon allocation during AM symbioses.

### Arbuscular mycorrhiza-induced changes in the expression of host abiotic and biotic stress response genes are species specific

Although the primary focus of this study was to evaluate both resource exchange and transporter expression, our gene expression data also provided some unique insights into biotic and abiotic stress responses. Mycorrhizal growth responses are context-dependent and fall along a mutualism to parasitism continuum (Johnson and Graham 2013). In natural environments, plants are colonized by communities of AM fungi, which supports the idea of functional complementarity, where a host plant experiences a broad range of benefits that are provided by specific members of the AM fungal community (*e.g.*, enhanced nutrient availability or increased disease resistance; Jansa et al., 2008). While Ga165 does not seem to provide sufficient nutritional benefits to elicit a positive growth response, it is possible that like other AM fungal species it may provide resistance to one or more abiotic stresses (*e.g.*, salinity or heavy metal accumulation), none of which were evaluated in this study.

We found that Ga165 more strongly up-regulated defense-response genes in the shoots (**Fig. 8B**), two of which (*Medtr4g120760* and *Medtr8g096900*) are upregulated in alfalfa when challenged with the foliar fungal pathogen *Phoma medicaginis* (Li et al. 2019a) and pea aphids (*Acyrthosiphon pisum*; Li et al., 2019b). There are two potential explanations for this observation. The plant could show similar responses to a low benefit AM fungus than to known fungal pathogens, or Ga165 could be more effective than Ri09 at priming host defense responses and thereby prevent infection from pathogens and mitigating arthropod attack as previously been described for other AM fungal species (Liu et al. 2007; Sharma et al. 2017).

Similarly, we also observed that genes associated with abiotic stress were upregulated in Ga165-colonized roots (**Supplemental Fig. S6A)**. Three of these genes — *Medtr5g063670*, *Medtr7g093170*, and *Medtr8g026960* — were very strongly upregulated and are annotated as annexin D8, seed maturation protein, and homeobox associated leucine zipper protein, respectively. Although the precise role of annexin D8 is not known, other annexins are differentially regulated during various abiotic and biotic stresses, and some seem to play a role during AM symbiosis (Roux et al. 2012). The seed maturation protein *Medtr7g093170* has been characterized as the late embryogenesis abundant protein LEA1370, which is upregulated under various abiotic stresses, including abscisic acid treatment, cold, salinity and dehydration (Zhang et al. 2020). Finally, the homeobox associated leucine zipper protein encoded by *Medtr8g026960* was characterized as a transcription factor that is upregulated during salt stress in alfalfa (Postnikova et al. 2013). However, since these genes are not yet functionally characterized, the question whether Ga165 colonization could contribute to an alleviation of these abiotic stresses, or whether host plants respond similarly to abiotic stress and to Ga165 colonization, requires further studies.

It is also possible that Ga165 truly is a low-benefit symbiont capable of persisting in a community with high benefit AM fungi thereby avoiding both detection and subsequent sanctioning by the host plant for poor performance (Hart et al. 2013). Avoiding detection could be carried out using effectors, as has been described in the model AM fungus *R. irregularis* DAOM197198 that counteracts the plant immune program with the small, secreted peptide SP7 (Kloppholz et al. 2011). Indeed, in our root expression data we observed that 11 defense response genes, including three defensins (*Medtr8g012810, Medtr8g012815*, and *Medtr8g012845*) were strongly up-regulated in Ri09, but not in Ga165-colonized roots (**Supplemental Fig. S6B**, **Supplemental Table S5**). Defensins are known to inhibit pathogen growth (Maróti et al. 2015), so it is tempting to speculate that Ga165 could temper host defense responses to avoid detection and sanctioning. However, the defensin *DefMd1* when overexpressed upregulates the expression of the GRAS transcription factor *RAM1*, which plays a critical role in arbuscular branching (Park et al., 2015; Uhe et al., 2018). *RAM1* was highly upregulated in both Ga165 and Ri09 colonized roots (**Supplemental Table S3**) suggesting that defensins might play a role in AM symbiosis.

### Transporter expression patterns in *Medicago truncatula* roots reveal interesting candidates for future functional characterization

This study has provided novel insights into the physiological and transcriptomic response of a host plant to colonization by both high and low benefit AM fungi and **Fig. 9** provides an overview of these responses. The experimental approach allowed us to identify both the molecular mechanisms controlling the observed physiological responses and transporters that are both commonly and uniquely regulated by high (Ri09) and low benefit (Ga165) AM fungal species. While the primary focus of our study was on N, P, and carbon transporters in the roots of *M. truncatula* colonized by Ga165 or Ri09, our analysis of differentially expressed transporters uncovered some interesting gene candidates with unique expression patterns (**Figs. 6** and **7**). Although there was a core set of transporters similarly induced by both fungi, a smaller set were expressed differently, and many are prime candidates for future functional analysis.

**Figure 9.**
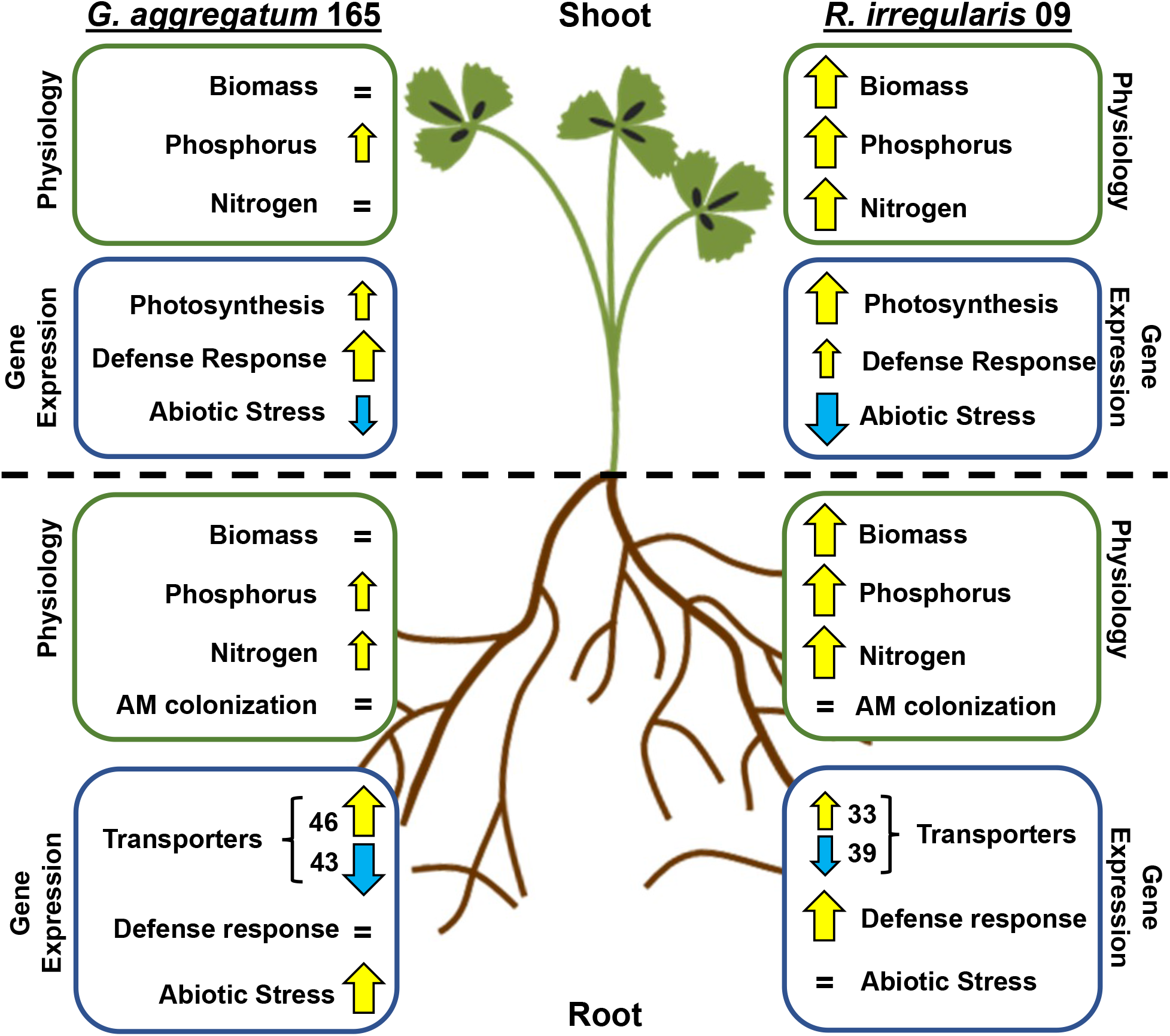
Summary of the physiological and transcriptomic responses in roots and shoots of *Medicago truncatula* colonized by either *Glomus aggregatum* 165 or *Rhizophagus irregularis* 09. Physiological data include the analysis of AM colonization, plant biomass, and both phosphorus and nitrogen-15 contents in plant tissues. Transcriptomic data include the expression patterns of nutrient transporters and enrichment of gene ontology terms. Yellow up-arrows indicate an increase, equal signs (=) specify no change, and blue down-arrows designate a decrease in each response compared to non-mycorrhizal plants. Arrow size indicates the degree of the response.

Multiple ABC transporters were induced by AM colonization (**Fig. 7C**), and two of these, *STR* and *STR2*, have already been studied extensively (Zhang et al., 2010). Two other transporters, *AMN2* and *AMN3*, are also well studied, but the substrate they transport remains elusive (Roy et al. 2021). The two remaining genes, *Medtr4g124040* and *Medtr1g050525*, have not been functionally characterized and could represent interesting targets for future studies. The amino acid transporters *Medtr1g030660* and *Medtr2g101920* and the peptide transporters *PTR3* and *OPT15* were all upregulated in mycorrhizal roots. None of these have been functionally characterized, but the protein product of *Medtr2g101920* was isolated from the plasma membrane of AM roots (Aloui et al. 2018). Future characterization of these transporters will provide greater insight into the extensive nutrient exchange mechanisms that support the AM symbiosis. In conclusion, the comparison of the physiological and transcriptomic response of *M. truncatula* to colonization with two AM fungal species that provide different nutritional benefits to the host allowed us to identify some of the molecular mechanisms that control reciprocal nutrient exchange. These include specific N and P transporters that have already been functionally characterized (e.g., *AMT2;3*, *AMT2;5*, *NPF4.12*, *PT4,* and *PT8*) and others that have not (e.g., NIP1;5). In addition, this study uncovered differential expression of genes putatively involved in the allocation of carbon resources to AM fungi depending on their nutritional benefit (e.g., *RAM2, SWEET1.2, SWEET3.3 and SWEET12* were upregulated in Ri09-colonized roots, while *SWEET7* and *SWEET13* were upregulated in Ga165-colonized roots). Root colonization with diverse fungal communities that are composed of both low and high benefit fungi, allows the host plant to choose among multiple trading partners, but also to take advantage of the diverse benefits that fungi within this community can provide. The differential gene expression in the roots and shoots after colonization with AM fungi that differ in the symbiotic benefit that they provide, indicates that host plants are able to fine tune their response to different fungi in their community.

## MATERIALS AND METHODS

### Biological materials and growth conditions

*M. truncatula* ‘Jemalong A17’ seeds were acid scarified in 36N H_2_SO_4_ (Fisher Scientific Inc., Waltham, MA, USA), surface sterilized with 8.25% sodium hypochlorite (Clorox^®^ bleach, The Clorox Company, Oakland, CA, USA), rinsed with sterile Type 1 water and imbibed at 4°C overnight. The seeds were then transferred onto moist autoclaved filter paper in Petri dishes and kept in the dark for 3 d. Finally, the Petri dishes were placed in ambient conditions for 7 d. Fully germinated seedlings were transferred into the root compartment (RC) of a custom-made, two-compartment pot measuring 12 cm x 8 cm x 8 cm (L x W x H; **Supplemental Fig. S1**). The RC was separated from the hyphal compartment (HC) by a 0.1 cm-thick plastic divider sealed on all sides by silicone (Aqueon, Franklin, WI, USA). A hole in the middle of the divider (~3.12 cm in diameter) was covered on both sides with a sheet of fine nylon mesh with 50 μm pores. Between these sheets was a coarse nylon mesh with 1000 μm pores to create an air gap and inhibit mass flow between the RC and the HC. This allowed fungal hyphae, but not roots, to crossover from the RC to the HC (**Supplemental Fig. S1**). Both compartments were filled with 250 ml of sterilized soil substrate containing 40% sand, 20% perlite, 20% vermiculite, and 20% soil by volume. The soil substrate had an original concentration of 4.81 mg kg^−1^ of Olsen’s extractable phosphate, 10 mg kg^−1^ of NH_4_^+^, 34.4 mg kg^−1^ of NO_3_^−^, and a pH of 8.26. These nutrient levels were sufficiently low to induce host demand for N and P and to stimulate AM colonization.

Fungal inoculum for *Rhizophagus irregularis* isolate 09 (Ri09; collected from Southwest Spain by Mycovitro S.L. Biotechnología ecológica, Granada, Spain) and *Glomus aggregatum* isolate 165 (Ga165; collected from the Long Term Mycorrhizal Research Site, University of Guelph, Canada) were produced using axenic root organ cultures of Ri T-DNA transformed carrot (*Daucus carota* clone DCI) grown on minimal medium (St-Arnaud et al. 1996). After approximately eight weeks of growth, spores were isolated by blending the medium in 10 mM citrate buffer (pH 6.0). Then, at transplanting, each seedling was inoculated with ~0.4 g mycorrhizal roots and ~500 spores of either Ri09 or Ga165. Non-mycorrhizal (NM) controls received a similar quantity of double autoclaved roots and spores. After transplanting and fungal inoculation, the plants were grown in a growth chamber (model TC30; Conviron, Winnipeg, MB, Canada) with a 25°C day/20°C night cycle, 30% relative humidity, and a photosynthetic photon flux of ~225 μmol m^−2^ s^−1^.

### Experimental design

Throughout the experiment, the position of the plants within the growth chamber was randomized three times. Five weeks post-inoculation, 4 mM ^15^NH_4_Cl (Sigma Aldrich, St. Louis, USA) and 0.5 mM KH2PO4 were added to the HC of colonized and noncolonized plants in a modified Ingestad solution (Ingestad 1960). The plants were harvested two weeks later (seven weeks post-inoculation).

### Biomass determination, mycorrhizal quantification, and analysis of nitrogen and phosphorus contents in plants

At plant harvest, shoots and roots were separated to determine their respective fresh weight. Subsamples of both tissues were flash frozen in liquid N2 and stored at −80°C for transcriptome analysis. An additional root subsample was collected and stored in 50% ethanol at 4°C to assess AM colonization. The remaining plant tissue was dried at 70°C for 48 h and then shoot and root dry weights were recorded. To determine the level of AM colonization, the root samples were cleared with 10% KOH at 80°C for 30 min, rinsed with tap water, and stained with 5% Sheaffer ink-vinegar (v:v) at 80°C for 15 min (Vierheilig et al. 1998). Total AM colonization rates were assessed using the gridline intersection method (McGonigle et al. 1990). We did not observe any fungal structures in the roots of NC control plants. Subsamples of dried shoot and root tissues were pulverized using a tissue homogenizer (Precellys 24 Dual, Cayman Chemical Company, Ann Arbor, MI, USA) and digested with 2N HCl for 2 h at 95°C. The P concentration in the plant tissues was spectrophotometrically determined at 436 nm after adding ammonium molybdate vanadate solution (Fisher Scientific, Pittsburgh, USA). Measurements of nitrogen-15 in plant tissues were performed by quantitative NMR spectroscopy as described previously (Kafle et al. 2018).

### Fluorescent staining and confocal analysis of arbuscules

Ink-stained roots that contained arbuscules were subsequently stained for confocal imaging as described in Cope et al. (2019). Briefly, they were incubated overnight at 4°C in 1X PBS containing 2 μg ml^−1^ Alexa Fluor 488™ conjugated to wheat germ agglutinin (WGA; Thermo Fischer Scientific, Waltham, MA, USA). The next day, stained roots were rinsed with 1X PBS, mounted on a glass slide, and immediately observed on an Olympus FV1200 confocal laser scanning microscope (Olympus, Shinjuku, Tokyo, Japan) with an Olympus UPlanApo 40×/0.85 objective. The emission spectra of Alexa Fluor 488™ WGA (500 to 540 nm) was detected after excitation with a 488-nm laser. Other settings used in the Fluoview v4.2c acquisition software (*e.g.*, laser intensity, gain, offset, magnification, airy units) were similar for all sample observations. The width of 488 and 508 arbuscules from 22 Ga165 and 21 Ri09-colonized root segments, respectively, were measured using Fiji (Schindelin et al. 2012).

### RNA sequencing and analysis

Root and shoot tissues from four biological replicates (n = 4) were homogenized in liquid N2, and total RNA was extracted using the PureLinkTM RNA Mini Kit (Thermo Fisher and quantified using a NanoDrop ND-2000 spectrophotometer (Thermo Fisher Scientific). RNA quality was assessed using 2100 BioAnalyzer technology (Agilent Technologies). Approximately 100 ng of total RNA was used to construct poly(A) selection libraries using the Illumina TruSeq RNA Sample Preparation kit and 150-nucleotide single-end reads were sequenced using an Illumina NextSeq500 with high output flow cells (Genomics Sequencing Facility, South Dakota State University).

The RNA sequencing data were uploaded to Discovery Environment (https://de.cyverse.org/de) and read quality of each fastq file was evaluated using FastQC 0.11.5 (https://www.bioinformatics.babraham.ac.uk/projects/fastqc/). One Ga165 and one NC root sample and one Ri09 shoot sample were discarded due to poor quality. The remaining root (n=3, Ga165 and NC; n=4, Ri09) and shoot (n=4, Ga165 and NC; n=3, Ri09) samples were aligned to the reference genome of *M. truncatula* (JCVI Mt4.0; Tang et al., 2014) using HISAT2.1 (Kim et al. 2019). Aligned reads were then assembled and merged using StringTie-1.3.3 (Pertea et al. 2015) and the *M. truncatula* JCVI 4.0v2 annotation file (Tang et al. 2014). Finally, Cuffdiff2.1.1 (Trapnell et al. 2013) was used to quantify transcript abundance. The output files from CuffDiff2.1.1 were visualized in R v3.6.2 (The R Foundation 2020) using the CummeRbund package (Goff et al. 2012). For further analyses, the differential gene expression output files from CuffDiff2.1.1 were imported into Microsoft Excel (Microsoft 365^®^) and sorted to identify significantly differentially expressed genes (DEGs) with a log2 fold-change > 2 and a false discovery rate-adjusted q-value < 0.05. All DEGs outside this threshold were not considered for further analysis. Venn diagrams of DEGs were generated using the Bioinformatics & Evolutionary Genomics Web tool (http://bioinformatics.psb.ugent.be/webtools/Venn/) and heatmaps of FPKM (Fragments Per Kilobase of transcript per Million mapped reads) values from all DEGs across inoculation regimes were generated using Cluster 3.0 and Java TreeView (http://bonsai.hgc.jp/~mdehoon/software/cluster/software.htm). Gene ontology (GO) enrichment analysis was conducted with the web-based tool PlantTFDB 4.0 (http://plantregmap.gao-lab.org/go.php).

### Statistical analysis

For the physiological response variables, unless mentioned otherwise, mean and standard error of the mean were determined from five independent biological replicates for each colonization regime, including noncolonized and either Ga165 or Ri09-colonized plants. One-way ANOVA was used among all three colonization regimes for dry weight, P concentration, P content, and nitrogen-15 enrichment. The least significant difference (LSD) test was subsequently used for pairwise multipl treatments. These statistical analyses were conducted using Statistix 9 (analytical software, Tallahassee, FL, USA). For AM colonization and arbuscule size, statistically significant differences (p ≤ 0.05) between the Ga 165 and Rio9 mycorrhizal treatments were determined using a Welch’s two-sample t-test in R v4.0.2 (The R Foundation 2020). Only results that are significant at the p ≤ 0.05-level are primarily discussed. All F statistics and p-values are shown in **Supplemental Table S1**.

## ACKNOWLEDGEMENTS

This work was funded by the USDA (2017-67014-26530), the SD Soybean Research and Promotion Council, and the Agricultural Experiment Station at SDSU to HK and SS. KG acknowledges support of the North Carolina Agriculture Research Service (NCARS), the North Carolina Soybean Producers Association (2019-1656), and the USDA (2020-67013-31800). We would like to thank Lindsay McKeever for performing the Kjeldahl degradations for the ^15^N/^14^N analyses.

## SUPPORTING INFORMATION

**Supplemental Figure S1.** Schematic model of the dual-compartment pot system used for this study.

**Supplemental Figure S2**. Shoot and root phosphorous concentration of *Medicago truncatula* colonized by arbuscular mycorrhizal fungi.

**Supplemental Figure S3**. Hierarchical clustering of biological replicates from root RNA sequencing data.

**Supplemental Figure S4**. Hierarchical clustering of biological replicates from shoot RNA sequencing data.

**Supplemental Figure S5**. Volcano plots of differentially expressed genes (DEGs) between treatments.

**Supplemental Figure S6**. Summary of gene ontology enrichment analysis of genes upregulated in the roots of *Medicago trunctaula*.

**Supplemental Figure S7**. Summary of gene ontology enrichment analysis of genes downregulated in the roots of *Medicago trunctaula*.

**Supplemental Figure S8**. Alternative portrayal of expression patterns of nutrient transporter genes.

**Supplemental Figure S9**. Alternative portrayal of expression patterns of secondary metabolite transporter genes.

**Supplemental Figure S10**. Summary of gene ontology enrichment analysis of differentially expressed genes in the shoots of *Medicago truncatula*.

**Supplemental Figure S11**. Alternative portrayal of expression patterns of genes involved in photosynthesis, response to biotic stress, and response to abiotic stress.

**Supplemental Table S1.** F statistics and p-values from one-way ANOVA for each experimental parameter.

**Supplemental Table S2**. Expression levels of genes associated with the strigolactone biosynthesis pathway and the common symbiosis signaling pathway in roots of *Medicago truncatula*.

**Supplemental Table S3**. Expression levels of genes downstream of the common symbiosis signaling pathway in roots of *Medicago truncatula*.

**Supplemental Table S4**. List of differentially regulated genes from shoot gene ontology enrichment analysis.

**Supplemental Table S5**. List of differentially regulated defense response genes from root gene ontology enrichment analysis.

**Supplemental File 1**. Significantly differentially expressed genes in roots

**Supplemental File 2**. Significantly differentially expressed genes in shoots

**Supplemental File 3**. Significant gene ontology terms in roots

**Supplemental File 4**. Significantly differentially expressed transporters in roots

**Supplemental File 5**. List of mineral transporters by type

**Supplemental File 6**. List of secondary metabolite transporters by type

**Supplemental File 7**. Significant gene ontology terms in shoots

---

**Supplemental Figure S1.**
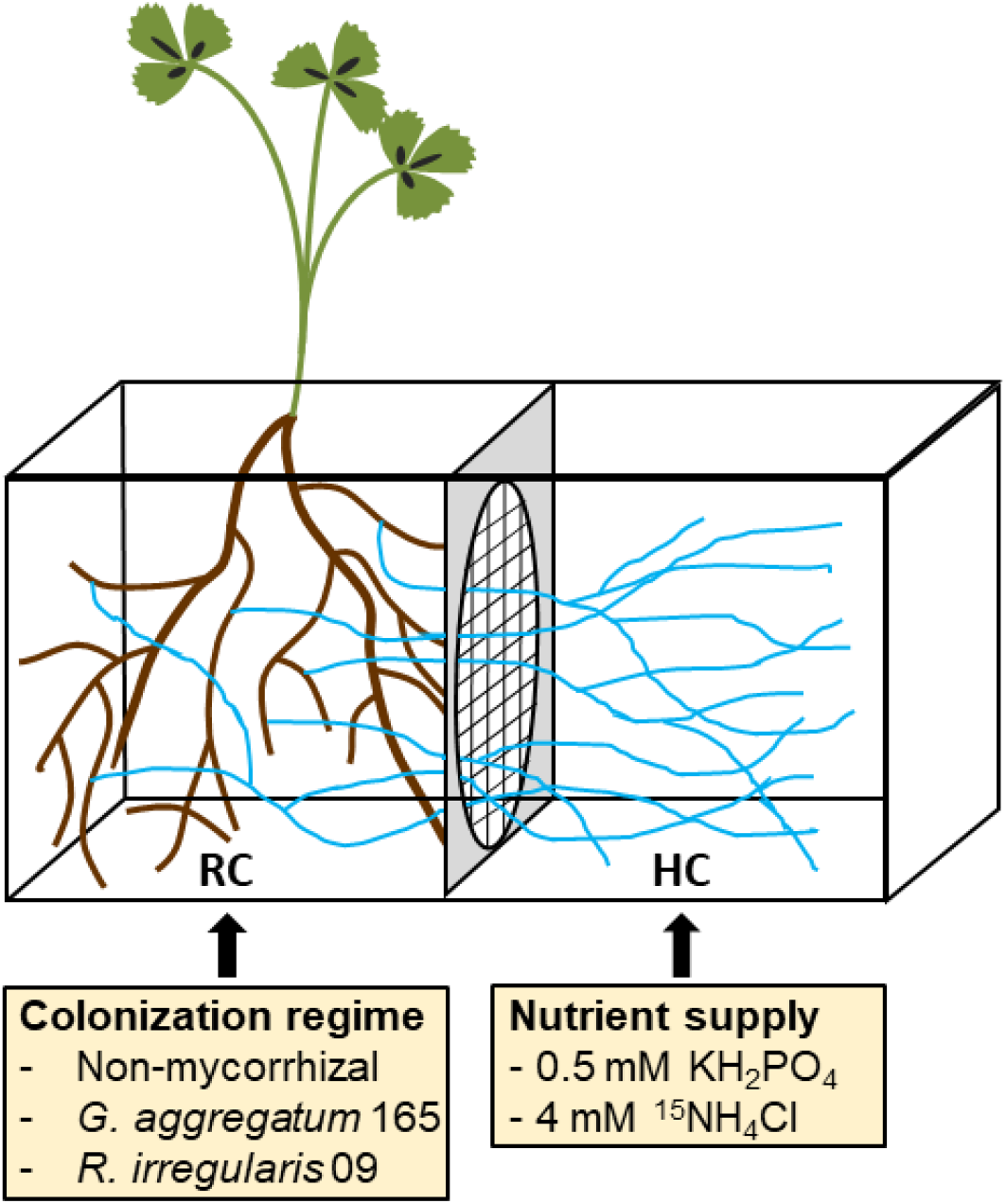
Schematic model of the dual-compartment pot system used for this study. Roots in the root compartment (RC) were mock-inoculated for the non-mycorrhizal control treatment and for the mycorrhizal treatments, roots were inoculated with either 500 spores of either *Glomus aggregatum* 165 or *Rhizophagus irregularis* 09. Five weeks post-inoculation, 0.5 mM KH_2_PO_4_ and 4 mM ^15^NH_4_Cl were added to the hyphal compartment (HC).

**Supplemental Figure S2.**
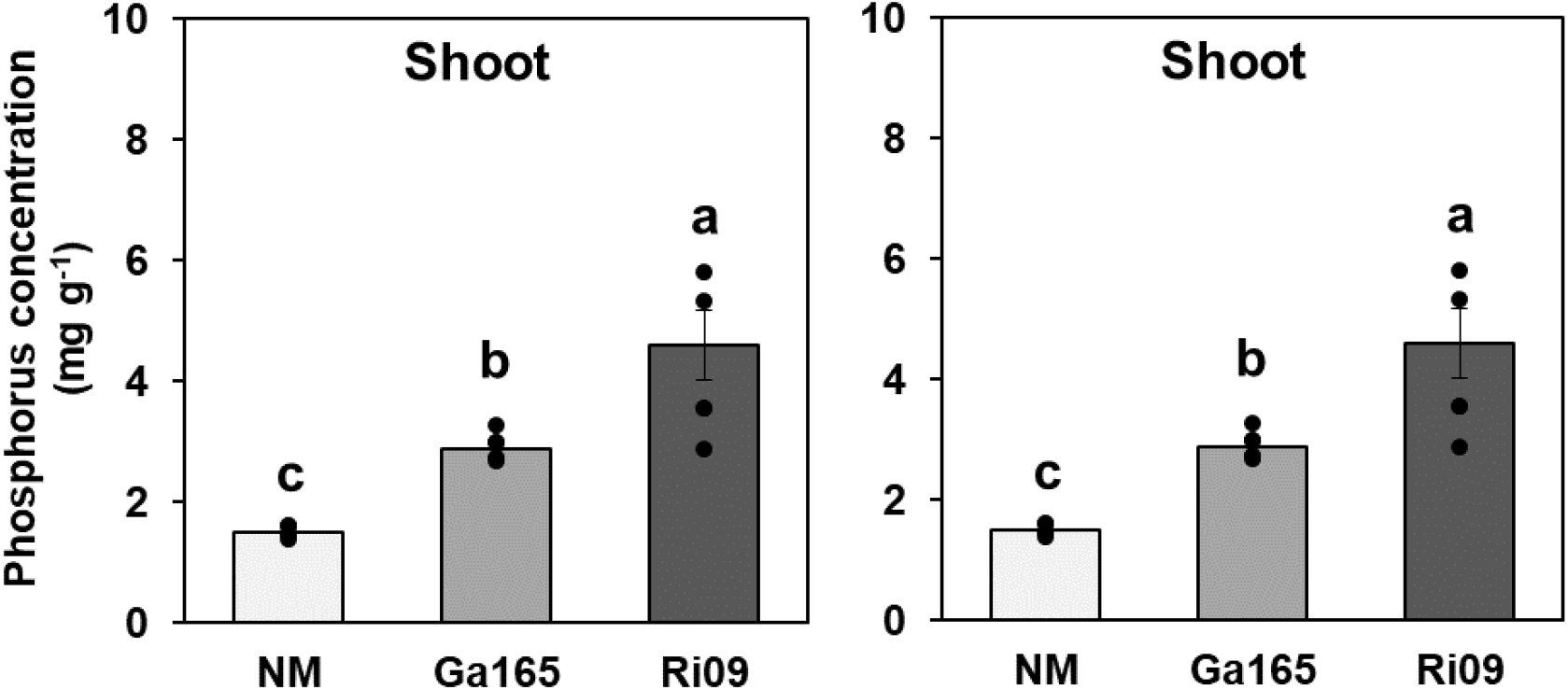
Shoot and root phosphorous concentration of *Medicago truncatula* colonized by arbuscular mycorrhizal fungi. The plants were inoculated with either *Glomus aggregatum* 165 (Ga165) or *Rhizophagus irregularis* 09 (Ri09). Control plants were mock inoculated and served as non-mycorrhizal (NM) controls. Data points represent individual values for biological replicates (n = 5) and bars represent the mean of each treatment ± SEM. Different letters above the bars indicate statistically significant differences (p ≤ 0.05) based on one-way ANOVA and LSD test. F statistics and p-values are shown in **Supplemental Table S1**.

**Supplemental Figure S3.**
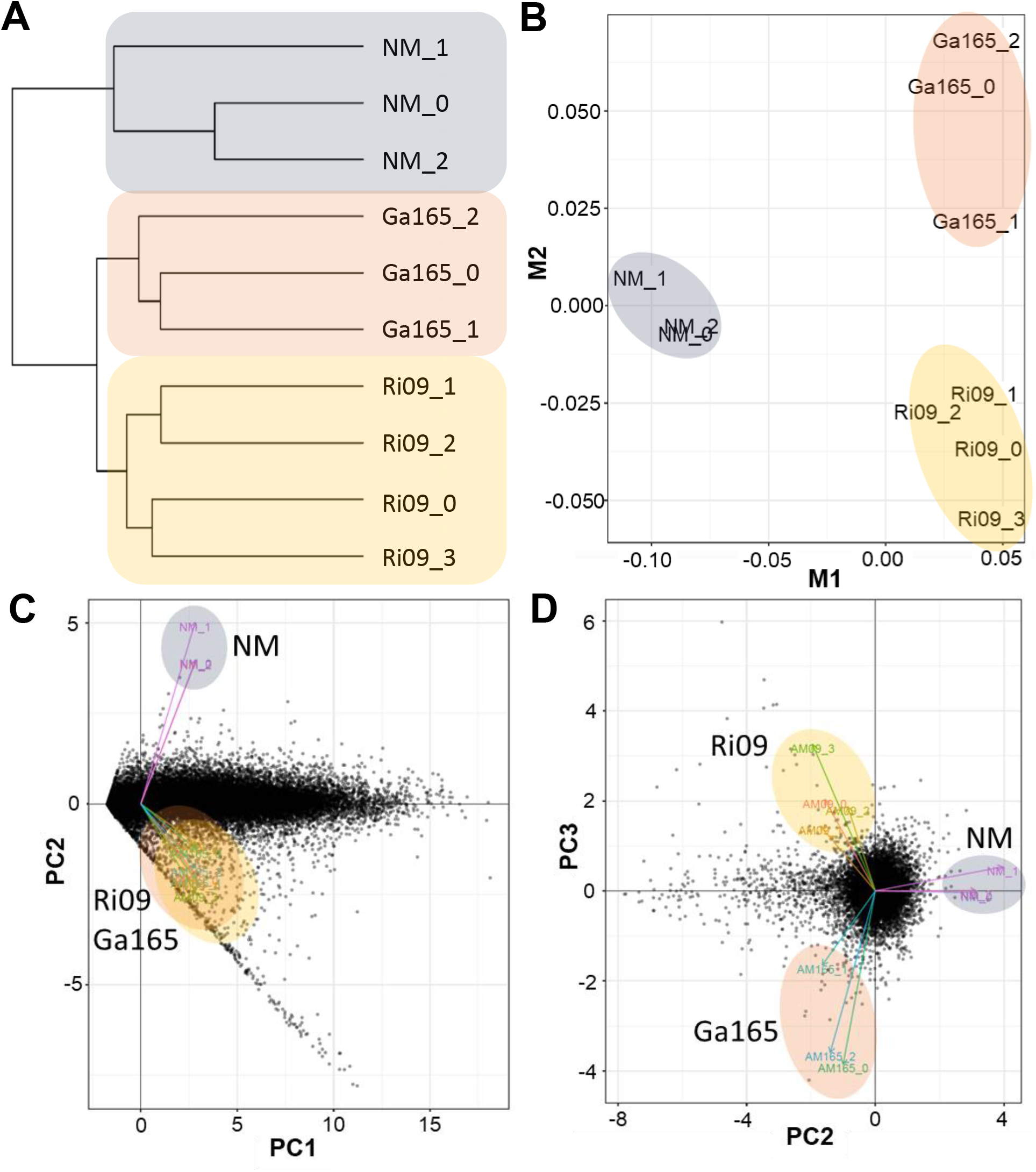
Hierarchical clustering of biological replicates from root RNA sequencing data. Treatments include non-mycorrhizal (NM, n = 3), *Glomus aggregatum* 165 (Ga165, n = 3) and *Rhizophagus irregularis* 09 (Ri09, n = 4). The data were clustered using **A**, dendrogram analysis; **B**, multidimensional scaling; and principal component analyses (PCA), including PC1 vs. PC2 (**C**) and PC2 vs. PC3 (**D**). Note the distinct clustering of all biological replicates from each treatment.

**Supplemental Figure S4.**
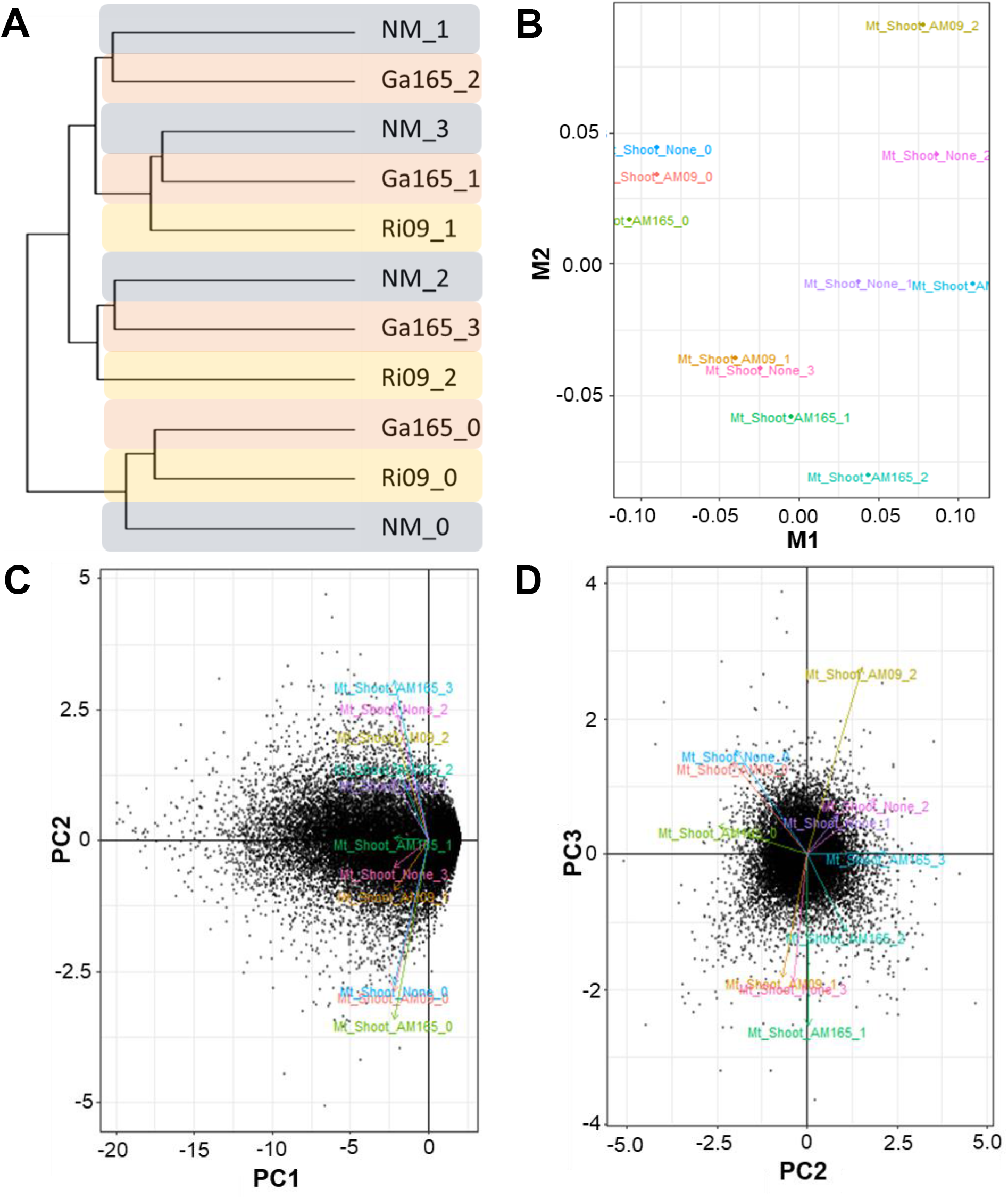
Hierarchical clustering of biological replicates from shoot RNA sequencing data. Treatments include non-mycorrhizal (NM, n = 4), *Glomus aggregatum* 165 (Ga165, n = 4) and *Rhizophagus irregularis* 09 (Ri09, n = 3). The data were clustered using **A**, dendrogram analysis; **B**, multidimensional scaling; and **C-D**, principal component analyses (PCA), including PC1 vs. PC2 (**C**) and PC2 vs. PC3 (**D**). Note that no distinct clustering of biological replicates was observed for any of the treatments.

**Supplemental Figure S5.**
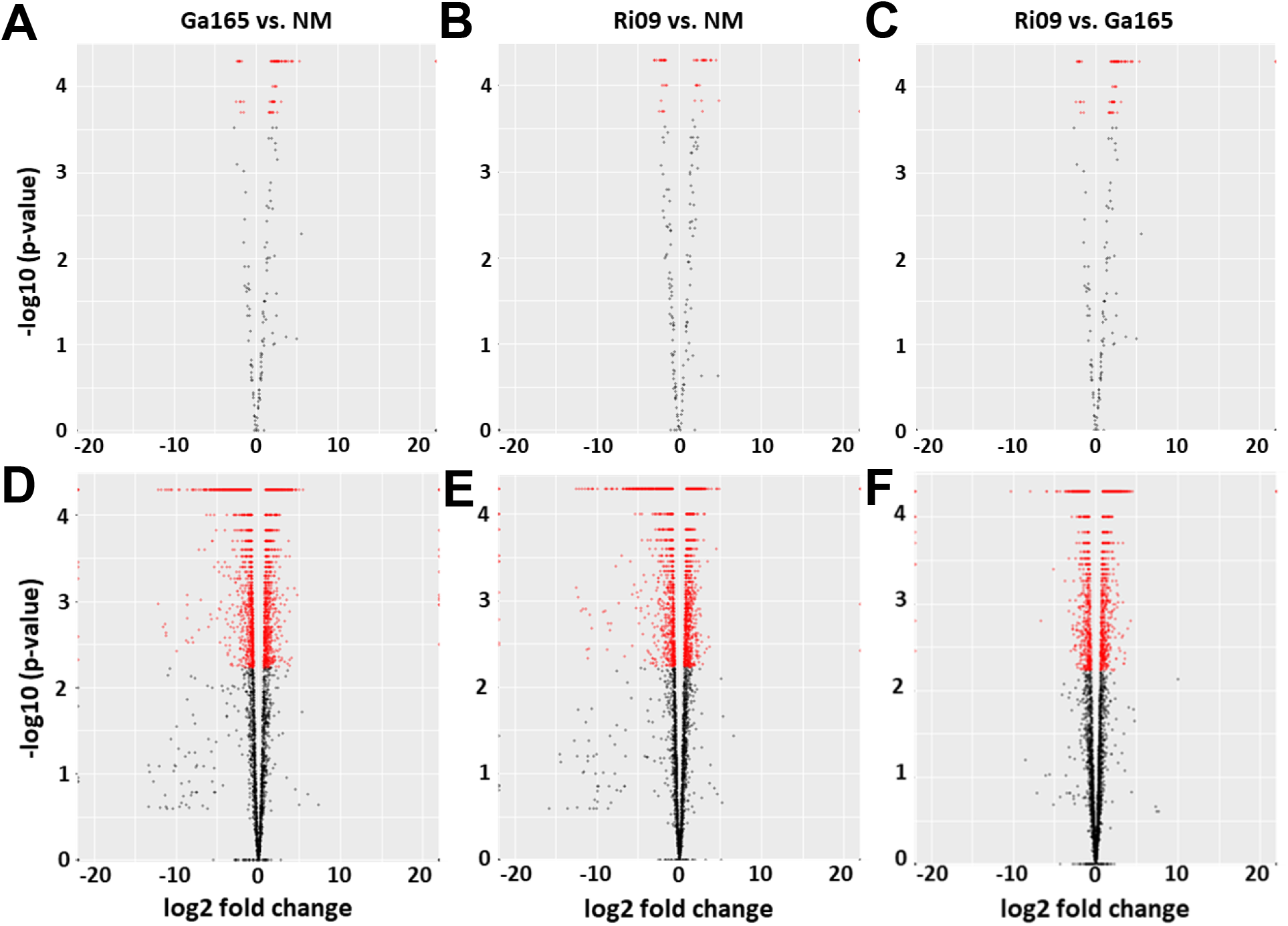
Volcano plots of differentially expressed genes (DEGs) between treatments. **A-C**, shoot data are shown on top and **D-F**, root data on bottom for the following comparisons: left, *Glomus aggregatum* 165 (Ga165) vs. non-mycorrhizal (NM); middle, *Rhizophagus irregularis* 09 (Ri09) vs. NM; and right, Ri09 vs. Ga165. Data points representing significant DEGs (-log_10_[p-value] > 2.3) are shown in red while non-significant DEGs (-log_10_[p-value] < 2.3) are shown in black. Note the greater abundance of significant DEGs among the root data compared to the shoot data.

**Supplemental Figure S6.**
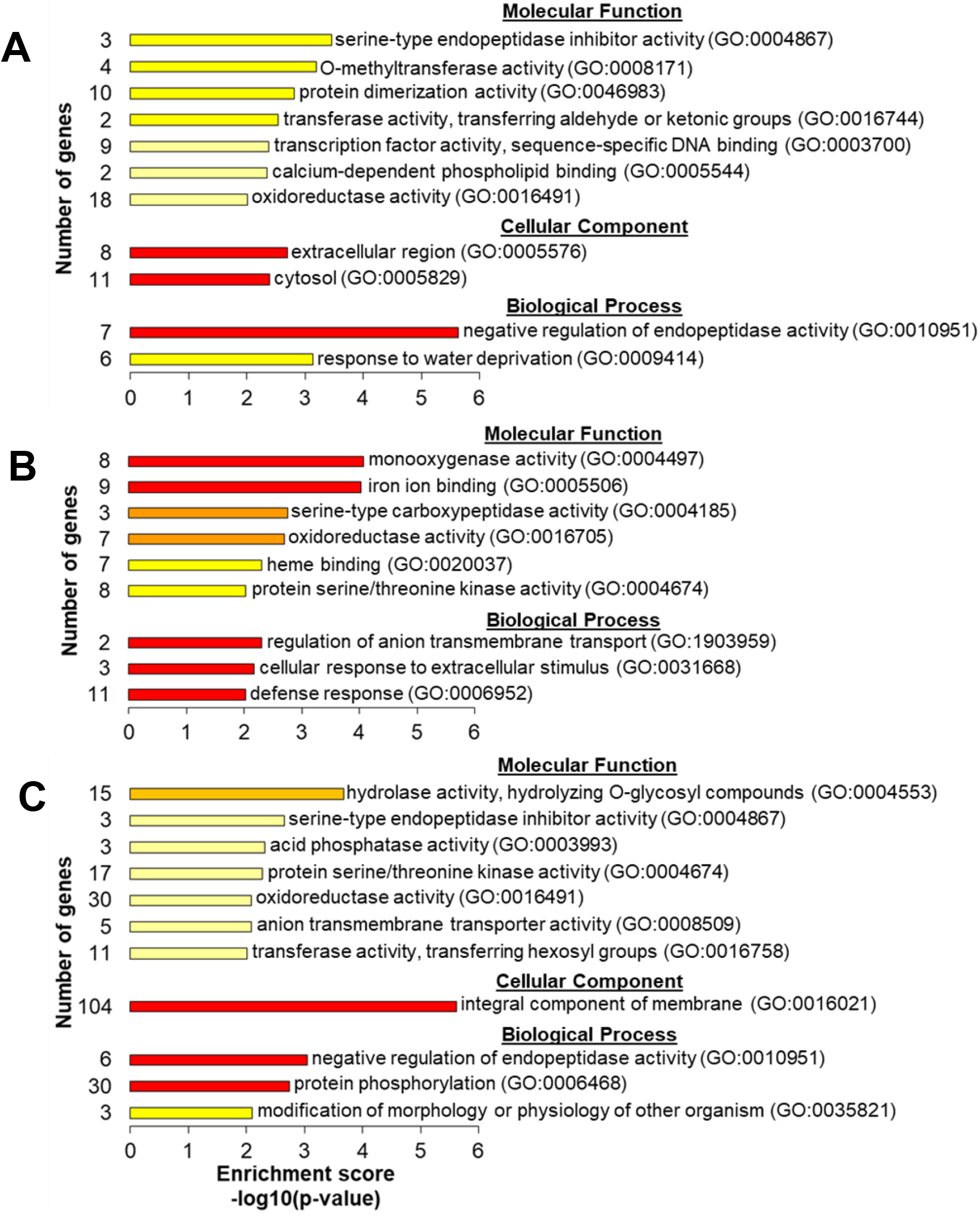
Summary of gene ontology enrichment analysis of genes upregulated in the roots of *Medicago truncatula*. All enriched gene ontology (GO) terms for **A**, *Glomus aggregatum* 165; **B**, *Rhizophagus irregularis* 09; or **C**, both species are shown for molecular function, cellular component, and biological process. The enrichment score (-log_10_[p-value]) for each GO term is also shown ranging from low (light yellow) to high (red). A list of the genes from each GO term can be found in **Supplemental File 4**.

**Supplemental Figure S7.**
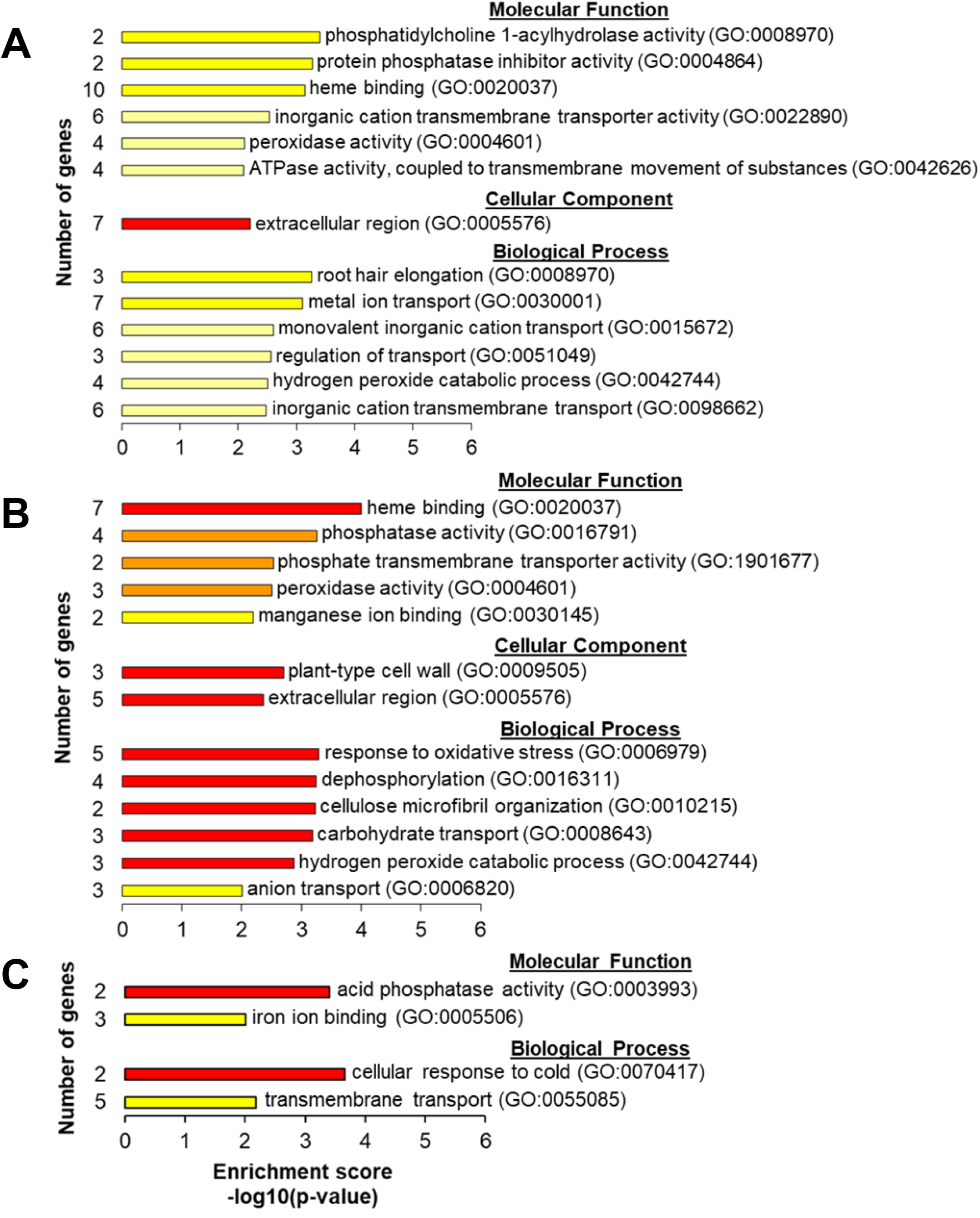
Summary of gene ontology enrichment analysis of genes downregulated in the roots of *Medicago trunctaula*. All enriched gene ontology (GO) terms for **A**, *Glomus aggregatum* 165; **B**, *Rhizophagus irregularis* 09; or **C**, both species are shown for molecular function, cellular component, and biological process. The enrichment score (-log_10_[p-value]) for each GO term is also shown ranging from low (light yellow) to high (red). A list of the genes from each GO term can be found in **Supplemental File 4**.

**Supplemental Figure S8.**
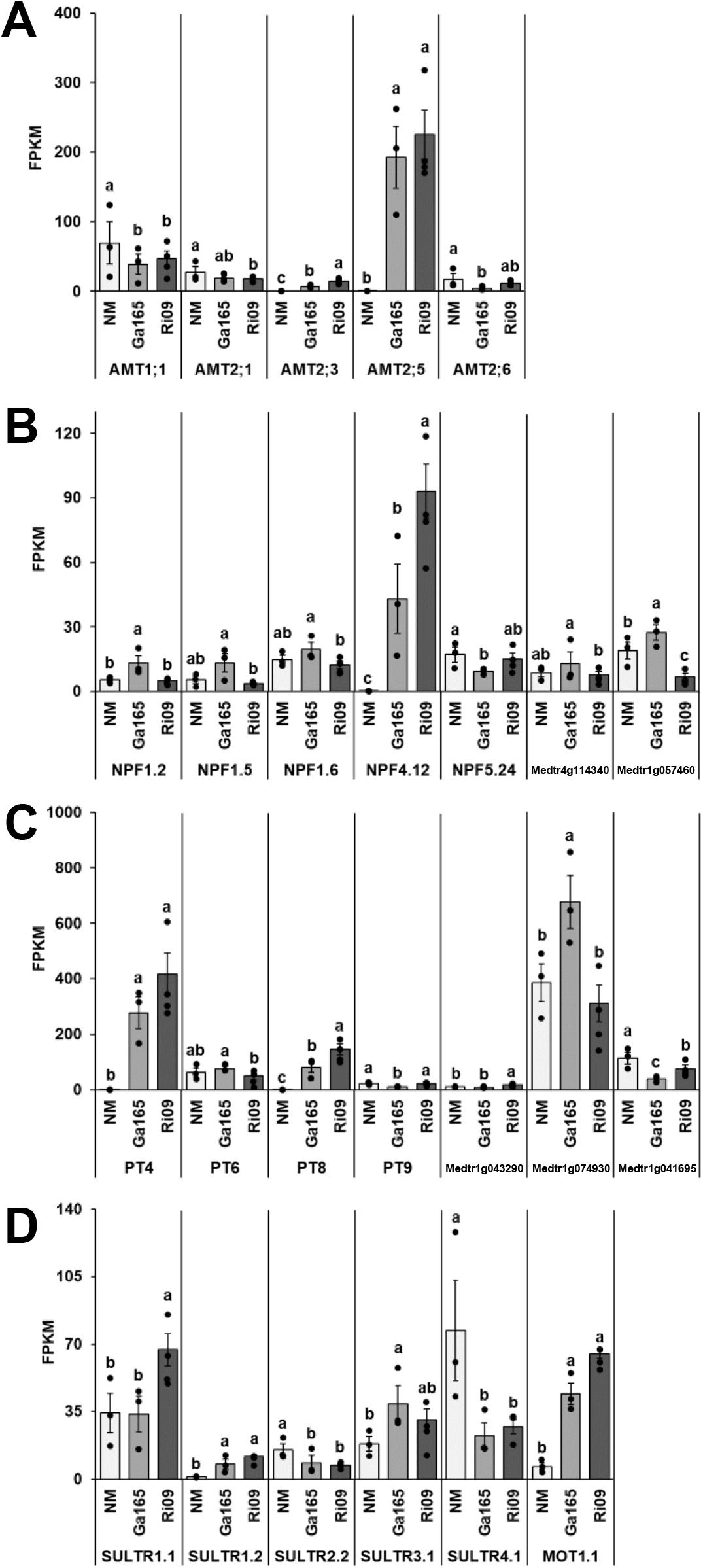
Alternative portrayal of expression patterns of genes annotated as **A**, ammonium; **B**, nitrate; **C**, phosphate; and **D**, sulfate transporters. Shown are individual data points for gene expression values from each biological replicate overlayed on the graphs from **Figure 6**, panels C through F, respectively.

**Supplemental Figure S9.**
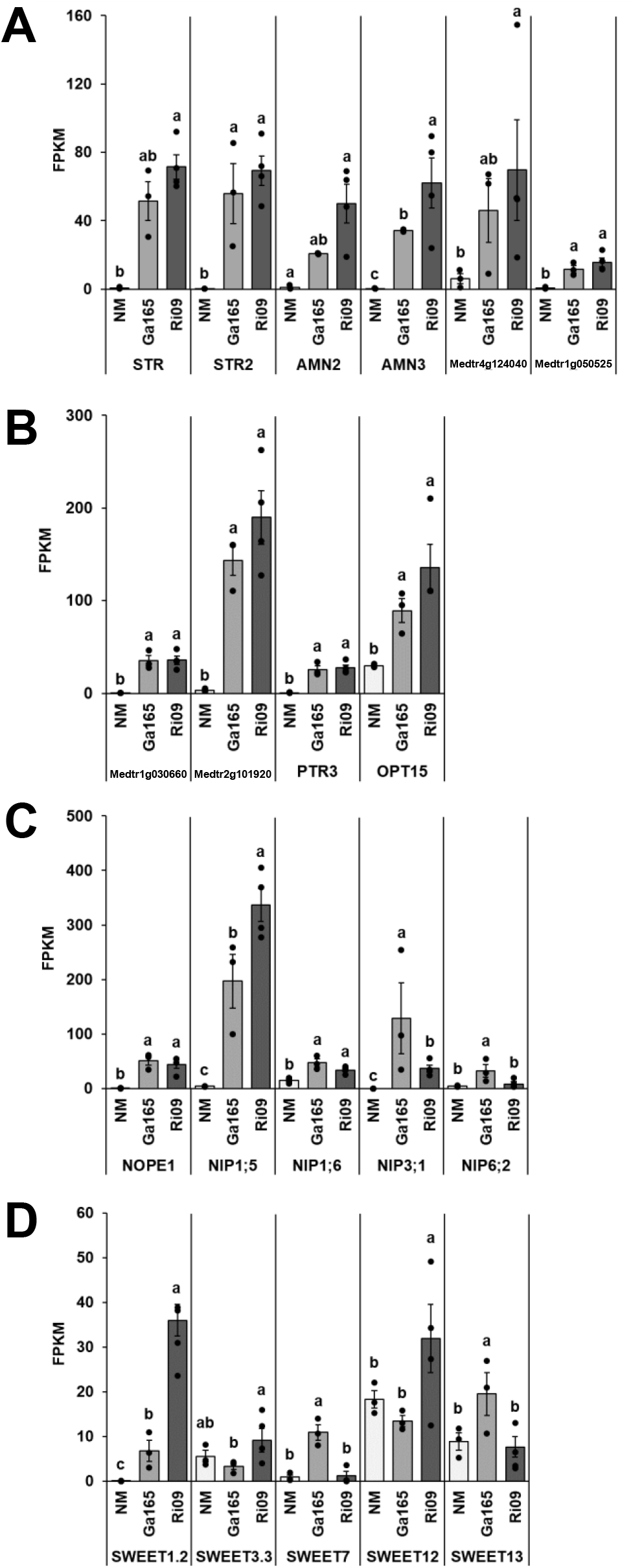
Alternative portrayal of expression patterns of genes annotated as **A**, ABC; **B**, amino acid or peptide; **C**, MIP or MFS; and **D**, SWEET transporters. Shown are individual data points for gene expression values from each biological replicate overlayed on the graphs from **Figure 7**, panels C through F, respectively.

**Supplemental Figure S10.**
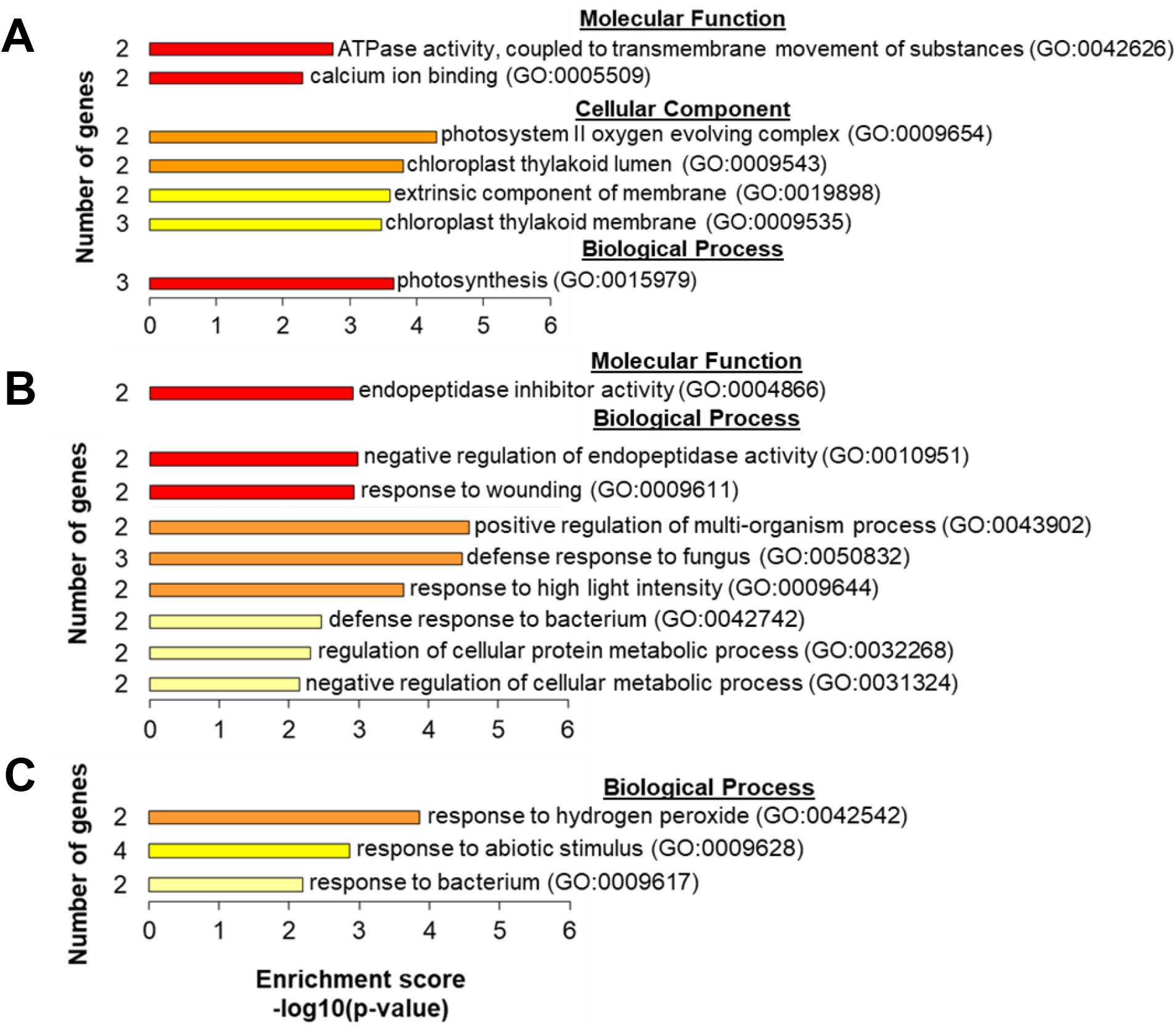
Summary of gene ontology enrichment analysis of differentially expressed genes in the shoots of *Medicago trunctaula*. Enriched gene ontology (GO) terms for molecular function, cellular component, and biological process among the following groups of genes based on the heatmap in **Figure 2F**: **A**, genes upregulated in the shoots of plants colonized by *Rhizophagus irregularis* 09 (Ri09); **B**, genes upregulated in the shoots of plants colonized by *Glomus aggregatum* 165 (Ga165); and **C**, genes downregulated in the shoots of plants colonized by Ri09. The enrichment score (-log_10_[p-value]) for each GO term is also shown ranging from low (light yellow) to high (red). A list of the genes from each GO term can be found in **Supplemental File 8**.

**Supplemental Figure S11.**
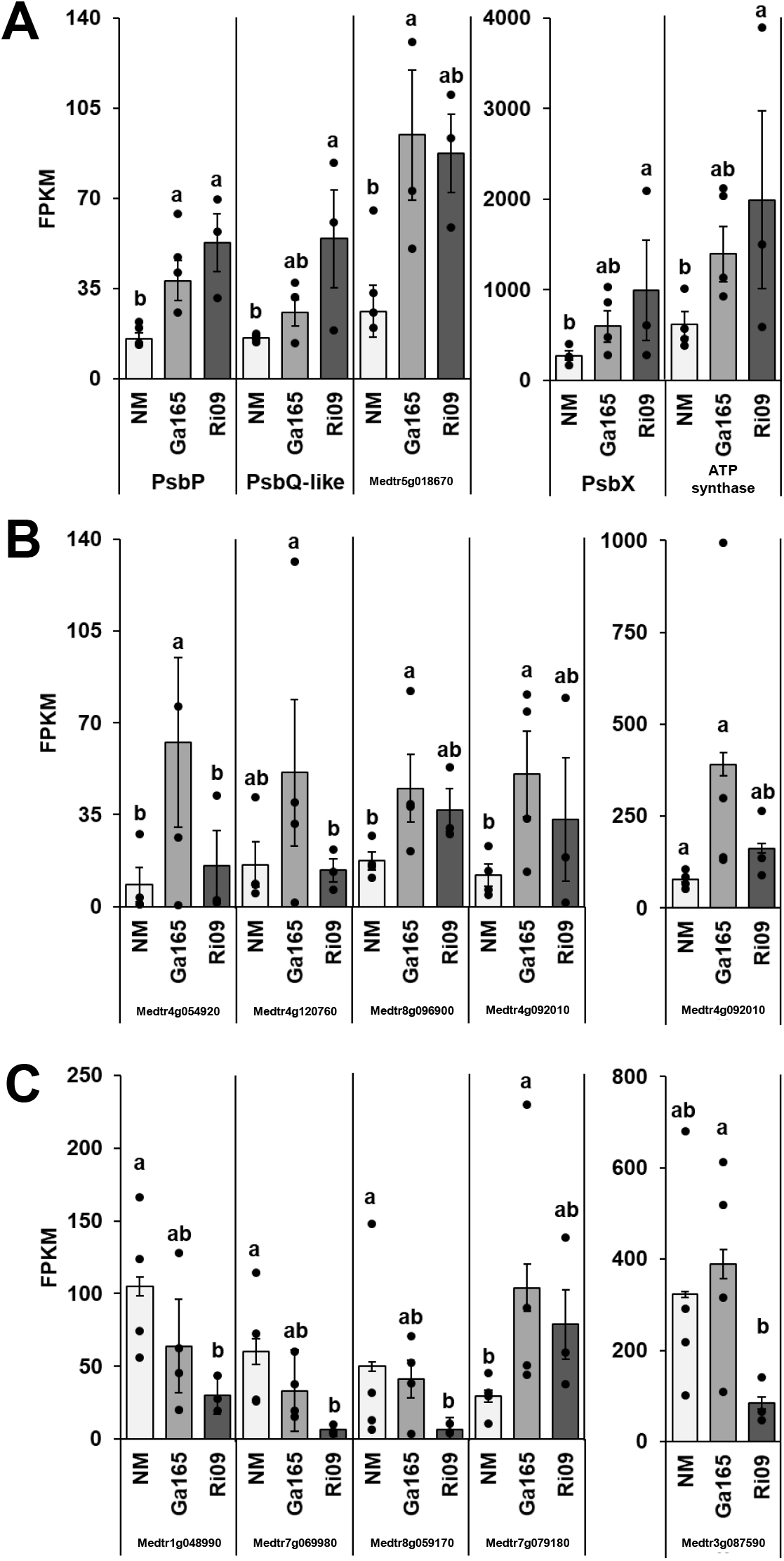
Alternative portrayal of expression patterns of genes involved in **A**, photosynthesis; **B**, response to biotic stress; and **C**, response to abiotic stress. Shown are individual data points for gene expression values from each biological replicate overlayed on the graphs from **Figure 8**, panels A through B, respectively.

**Supplemental Table S1.**
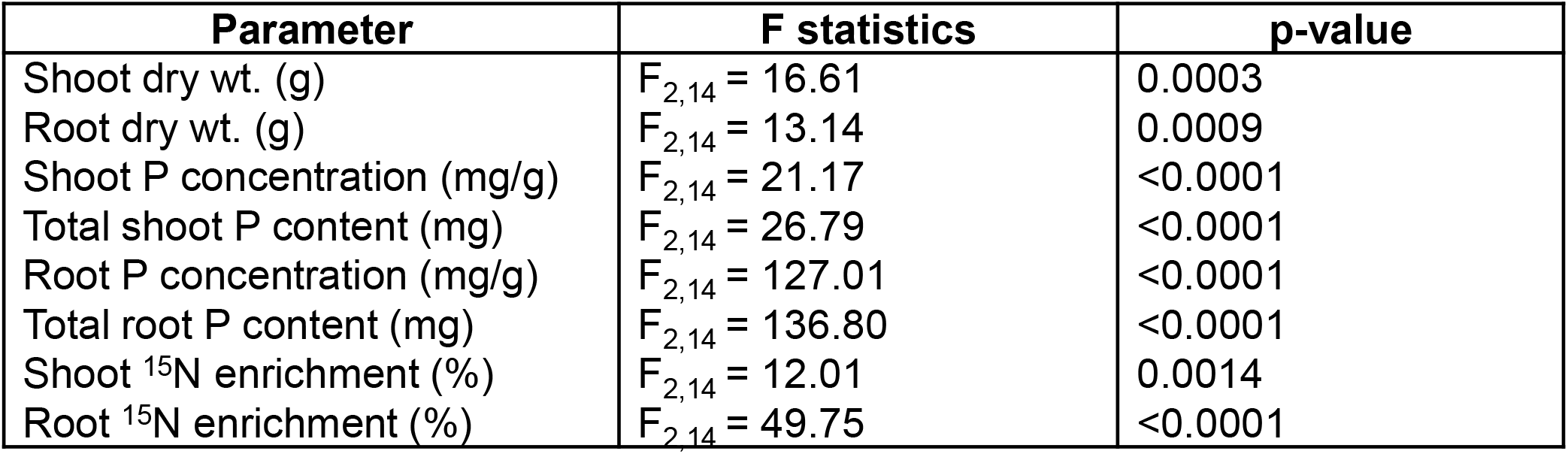
F statistics and p-values from one-way ANOVA for each experimental parameter.

| Parameter | F statistics | p-value |
| --- | --- | --- |
| Shoot dry wt. (g) | $F_{2,14} = 16.61$ | 0.0003 |
| Root dry wt. (g) | $F_{2,14} = 13.14$ | 0.0009 |
| Shoot P concentration (mg/g) | $F_{2,14} = 21.17$ | <0.0001 |
| Total shoot P content (mg) | $F_{2,14} = 26.79$ | <0.0001 |
| Root P concentration (mg/g) | $F_{2,14} = 127.01$ | <0.0001 |
| Total root P content (mg) | $F_{2,14} = 136.80$ | <0.0001 |
| Shoot $^{15}\text{N}$ enrichment (%) | $F_{2,14} = 12.01$ | 0.0014 |
| Root $^{15}\text{N}$ enrichment (%) | $F_{2,14} = 49.75$ | <0.0001 |

**Supplemental Table S2.**
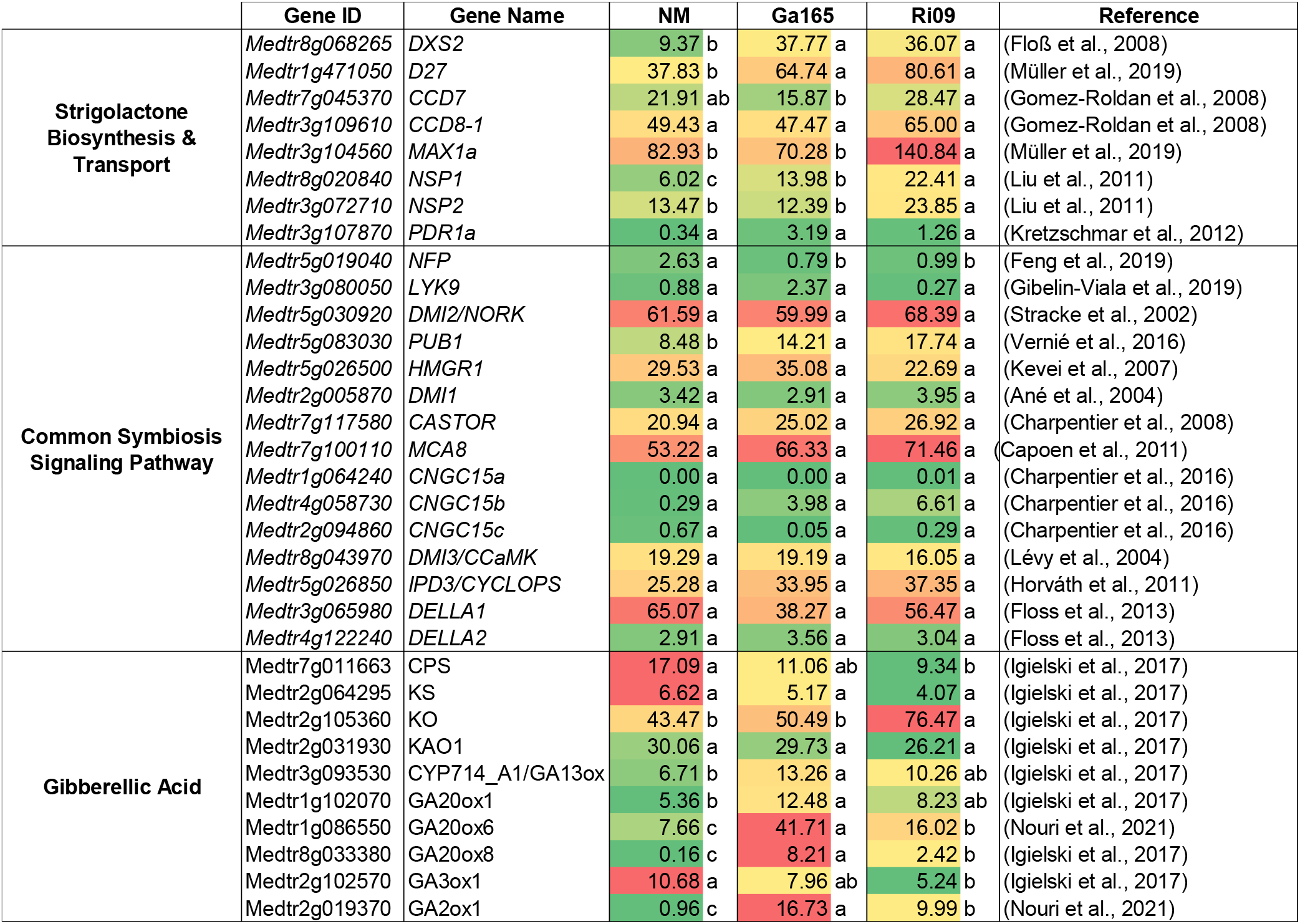
Expression levels of genes associated with the strigolactone biosynthesis pathway, the common symbiosis signaling pathway, and the gibberellic biosynthesis and degradation pathway in roots of *Medicago truncatula*. Mean FPKM (Fragments Per Kilobase of transcript per Million mapped reads) values for all listed genes are shown for all three treatments from this study, including nonmycorrhizal (NM, n=3) control plants and both *Glomus aggregatum* 165 (Ga165, n=3) and *Rhizophagus irregularis* 09 (Ri09, n=4) mycorrhizal plants. A green (low) to red (high) color scale is used to highlight differences in FPKM values. Significant differences (q-value < 0.05) in FPKM values between treatments were determined as part of the CuffDiff2 analysis and are indicated using different letters. References for the studies in which each gene was characterized are also included. Full citations can be found in the **Supplemental References**.

**Supplemental Table S3.** Expression levels of genes downstream of the common symbiosis signaling pathway in roots of *Medicago truncatula*. Mean FPKM (Fragments Per Kilobase of transcript per Million mapped reads) values for all listed genes are shown for all three treatments from this study, including nonmycorrhizal (NM, n=3) control plants and both *Glomus aggregatum* 165 (Ga165, n=3) and *Rhizophagus irregularis* 09 (Ri09, n=4) mycorrhizal plants. A green (low) to red (high) color scale is used to highlight differences in FPKM values. Significant differences (q-value < 0.05) in FPKM values between treatments were determined as part of the CuffDiff2 analysis and are indicated using different letters. References for the studies in which each gene was identified as playing a role in the arbuscular mycorrhizal symbiosis are also included. Full citations can be found in the **Supplemental References**.

|  | Gene ID | Gene Name | NM | Ga165 | Ri09 | Reference |
| --- | --- | --- | --- | --- | --- | --- |
| **Transcription Factors** | Medtr7g027190 | RAM1 | 0.01 b | 45.05 a | 52.08 a | (Park et al., 2015) |
|  | Medtr4g104020 | RAD1 | 0.09 c | 84.69 b | 139.93 a | (Park et al., 2015) |
|  | Medtr7g088990 | DIP1 | 111.61 a | 145.81 a | 137.94 a | (Yu et al., 2014) |
|  | Medtr2g034280 | MIG1 | 0.56 a | 0.97 a | 1.15 a | (Heck et al., 2016) |
|  | Medtr2g034260 | MIG2 | 9.07 b | 20.31 a | 18.69 a | (Heck et al., 2016) |
|  | Medtr2g034250 | MIG3 | 7.73 b | 18.70 a | 25.75 a | (Heck et al., 2016) |
|  | Medtr7g068600 | MYB1 | 0.06 c | 594.35 b | 1262.52 a | (Floss et al., 2017) |
| **Cellular Remodeling** | Medtr6g027840 | VAPYRIN/VPY | 3.20 b | 34.54 a | 33.33 a | (Pumplin et al., 2010) |
|  | Medtr1g017910 | EXO70I | 2.53 b | 24.18 a | 18.44 a | (Zhang et al., 2015) |
|  | Medtr2g088700 | SYP132A | 56.60 b | 92.13 a | 79.90 ab | (Pan et al., 2016) |
| **Lipid Biosynthesis & Transport** | Medtr8g468920 | WRI5a | 0.00 b | 2.26 a | 3.15 a | (Jiang et al., 2018) |
|  | Medtr8g107450 | STR | 0.68 b | 51.53 b | 71.44 a | (Zhang et al., 2010) |
|  | Medtr5g030910 | STR2 | 0.22 b | 55.77 a | 69.28 a | (Zhang et al., 2010) |
|  | Medtr1g040500 | RAM2 | 0.23 c | 192.67 b | 353.74 a | (Luginbuehl et al., 2017) |
|  | Medtr1g109110 | FatM | 0.07 a | 12.57 a | 17.47 a | (Bravo et al., 2017) |
| **Mineral Transport** | Medtr1g028600 | PT4 | 0.09 b | 277.24 a | 381.65 a | (Javot et al., 2007) |
|  | Medtr5g068140 | PT8 | 0.05 c | 81.31 b | 133.25 a | (Breuillin-Sessoms et al., 2015) |
|  | Medtr8g074750 | AMT2;3 | 0.00 c | 7.08 b | 13.07 a | (Breuillin-Sessoms et al., 2015) |
|  | Medtr7g115050 | AMT2;4 | 0.15 b | 6.53 a | 3.59 a | (Breuillin-Sessoms et al., 2015) |
|  | Medtr1g036410 | AMT2;5 | 0.11 b | 192.75 a | 213.55 a | (Breuillin-Sessoms et al., 2015) |
| **Sugar Transport** | Medtr3g089125 | SWEET1.2 | 0.04 c | 6.81 b | 32.93 a | (An et al., 2019) |

**Supplemental Table S4.** List of differentially regulated genes from shoot gene ontology enrichment analysis. Mean FPKM (Fragments Per Kilobase of transcript per Million mapped reads) values for all listed genes are shown for all three treatments from this study, including nonmycorrhizal (NM, n=3) control plants and both *Glomus aggregatum* 165 (Ga165, n=3) and *Rhizophagus irregularis* 09 (Ri09, n=4) mycorrhizal plants. A green (low) to red (high) color scale is used to highlight differences in FPKM values. Significant differences (q-value < 0.05) in FPKM values between treatments were determined as part of the CuffDiff2 analysis and are indicated using different letters.

|  | Gene ID | Gene Annotation | NM | Ga165 | Ri09 |
| --- | --- | --- | --- | --- | --- |
| **Photosynthesis** | *Medtr1g015290* | ultraviolet-B-repressible protein | 268.2 b | 662.9 ab | 992.1 a |
|  | *Medtr2g082580* | oxygen-evolving enhancer protein | 15.9 b | 28.6 ab | 54.4 a |
|  | *Medtr1g115410* | photosystem II reaction center PsbP family protein | 17.3 b | 44.6 a | 52.8 a |
|  | *Medtr1588s0010* | ATP synthase F1, gamma subunit | 605.3 b | 1556.0 ab | 1993.1 a |
|  | *Medtr6g084320* | ABC transporter-like family-protein | 12.8 b | 21.8 ab | 47.0 a |
| **Biotic Stress** | *Medtr4g054920* | cytochrome P450 family 94 protein | 8.26 b | 62.56 a | 15.45 b |
|  | *Medtr4g120760* | pathogenesis-related protein bet V I family protein | 16.02 ab | 51.03 a | 13.83 b |
|  | *Medtr5g053950* | allene oxide cyclase | 77.55 b | 391.10 a | 162.61 ab |
|  | *Medtr8g096900* | pathogenesis-related thaumatin family protein | 17.47 b | 45.09 a | 36.91 ab |
|  | *Medtr4g092010* | (3S)-linalool/(E)-nerolidol/(E,E)-geranyl linalool synthase | 11.90 b | 50.43 a | 33.24 ab |
| **Abiotic Stress** | *Medtr1g048990* | superoxide dismutase | 105.08 a | 63.93 ab | 30.32 b |
|  | *Medtr3g087590* | myo-inositol 1-phosphate synthase | 322.70 ab | 388.70 a | 84.82 b |
|  | *Medtr7g069980* | ferritin | 60.16 a | 33.23 a | 6.36 b |
|  | *Medtr8g059170* | NAC transcription factor-like protein | 49.87 a | 41.35 a | 6.64 b |
|  | *Medtr5g018670* | photosystem II oxygen-evolving enhancer protein | 35.97 b | 103.56 a | 87.44 ab |
|  | *Medtr7g079180* | late embryogenesis abundant protein | 29.47 b | 103.91 a | 78.75 ab |

**Supplemental Table S5.** List of differentially regulated defense response genes from root gene ontology enrichment analysis. Mean FPKM (Fragments Per Kilobase of transcript per Million mapped reads) values for all listed genes are shown for all three treatments from this study, including nonmycorrhizal (NM, n=3) control plants and both *Glomus aggregatum* 165 (Ga165, n=3) and *Rhizophagus irregularis* 09 (Ri09, n=4) mycorrhizal plants. A green (low) to red (high) color scale is used to highlight differences in FPKM values. Significant differences (q-value < 0.05) in FPKM values between treatments were determined as part of the CuffDiff2 analysis and are indicated using different letters.

|  | Gene ID | Gene Annotation | NM | Ga165 | Ri09 |
| --- | --- | --- | --- | --- | --- |
| Defense Response | *Medtr0012s0290* | disease resistance protein (TIR-NBS-LRR class), putative | 0.91 a | 3.53 b | 5.78 c |
|  | *Medtr1g015140* | WRKY family transcription factor | 0.67 a | 1.06 ab | 2.68 b |
|  | *Medtr2g008430* | LL-diaminopimelate aminotransferase | 2.08 a | 7.54 b | 14.17 c |
|  | *Medtr3g028040* | LRR and NB-ARC domain disease resistance protein | 0.08 a | 0.53 ab | 1.10 b |
|  | *Medtr3g072140* | disease resistance protein (TIR-NBS-LRR class), putative | 0.15 a | 0.56 ab | 0.82 b |
|  | *Medtr6g006990* | carbonic anhydrase family protein | 0.56 a | 3.53 ab | 7.55 b |
|  | *Medtr8g012810* | Defensin related | 0.00 a | 0.67 ab | 45.59 b |
|  | *Medtr8g012815* | Defensin related | 0.00 a | 1.00 ab | 34.68 b |
|  | *Medtr8g012845* | Defensin related | 0.00 a | 0.36 ab | 51.54 b |
|  | *Medtr8g012870* | Defensin related | 0.00 a | 0.49 ab | 5.59 b |
|  | *Medtr8g059405* | Defensin | 0.97 a | 0.73 a | 8.89 b |

